# Low coverage genomic data resolve the population divergence and gene flow history of an Australian rain forest fig wasp

**DOI:** 10.1101/2020.02.21.959205

**Authors:** Lisa Cooper, Lynsey Bunnefeld, Jack Hearn, James M Cook, Konrad Lohse, Graham N. Stone

## Abstract

Population divergence and gene flow are key processes in evolution and ecology. Model-based analysis of genome-wide datasets allows discrimination between alternative scenarios for these processes even in non-model taxa. We used two complementary approaches (one based on the blockwise site frequency spectrum (bSFS), the second on the Pairwise Sequentially Markovian Coalescent (PSMC)) to infer the divergence history of a fig wasp, *Pleistodontes nigriventris. Pleistodontes nigriventris* and its fig tree mutualist *Ficus watkinsiana* are restricted to rain forest patches along the eastern coast of Australia, and are separated into northern and southern populations by two dry forest corridors (the Burdekin and St. Lawrence Gaps). We generated whole genome sequence data for two haploid males per population and used the bSFS approach to infer the timing of divergence between northern and southern populations of *P. nigriventris,* and to discriminate between alternative isolation with migration (IM) and instantaneous admixture (ADM) models of post divergence gene flow. *Pleistodontes nigriventris* has low genetic diversity (π = 0.0008), to our knowledge one of the lowest estimates reported for a sexually reproducing arthropod. We find strongest support for an ADM model in which the two populations diverged *ca*. 196kya in the late Pleistocene, with almost 25% of northern lineages introduced from the south during an admixture event *ca.* 57kya. This divergence history is highly concordant with individual population demographies inferred from each pair of haploid males using PSMC. Our analysis illustrates the inferences possible with genome-level data for small population samples of tiny, non-model organisms and adds to a growing body of knowledge on the population structure of Australian rain forest taxa.

## Introduction

Division of an ancestral population into daughter populations is a universal and repeating process in biology. The tempo and mode of population divergence are central to evolutionary processes ranging from local adaptation and range expansion to the origin of species (Hey & Nielsen 2004; Martin et al., 2013; Sousa & Hey, 2013). From a demographic perspective, population divergence can be described in terms of the sizes of ancestral and descendant populations, the time at which the ancestral population split, and parameters capturing the timing, extent and direction of gene flow between descendant populations. Gene flow can be modelled in at least two general ways (Fig. 1): an isolation with migration (IM) model identifies gene flow as the result of ongoing dispersal (Nielsen & Wakeley 2001, Hey, & Nielsen 2004, Lohse et al 2011), while an instantaneous admixture (ADM) model associates gene flow with one or more discrete dispersal events in the past (Durand et al., 2011; Sousa & Hey, 2013; Lohse & Frantz, 2014). Thus, which of these models applies may tell us whether putative dispersal barriers are past or ongoing, and help to identify evolutionarily independent conservation units within species. The models also have very different implications for local adaptation: while local adaptation may proceed unimpeded during periods of complete isolation in the ADM model, continuous gene flow under the IM model imposes a constant genetic load of locally deleterious variants (Bisschop et al 2020).

**Figure 1.**
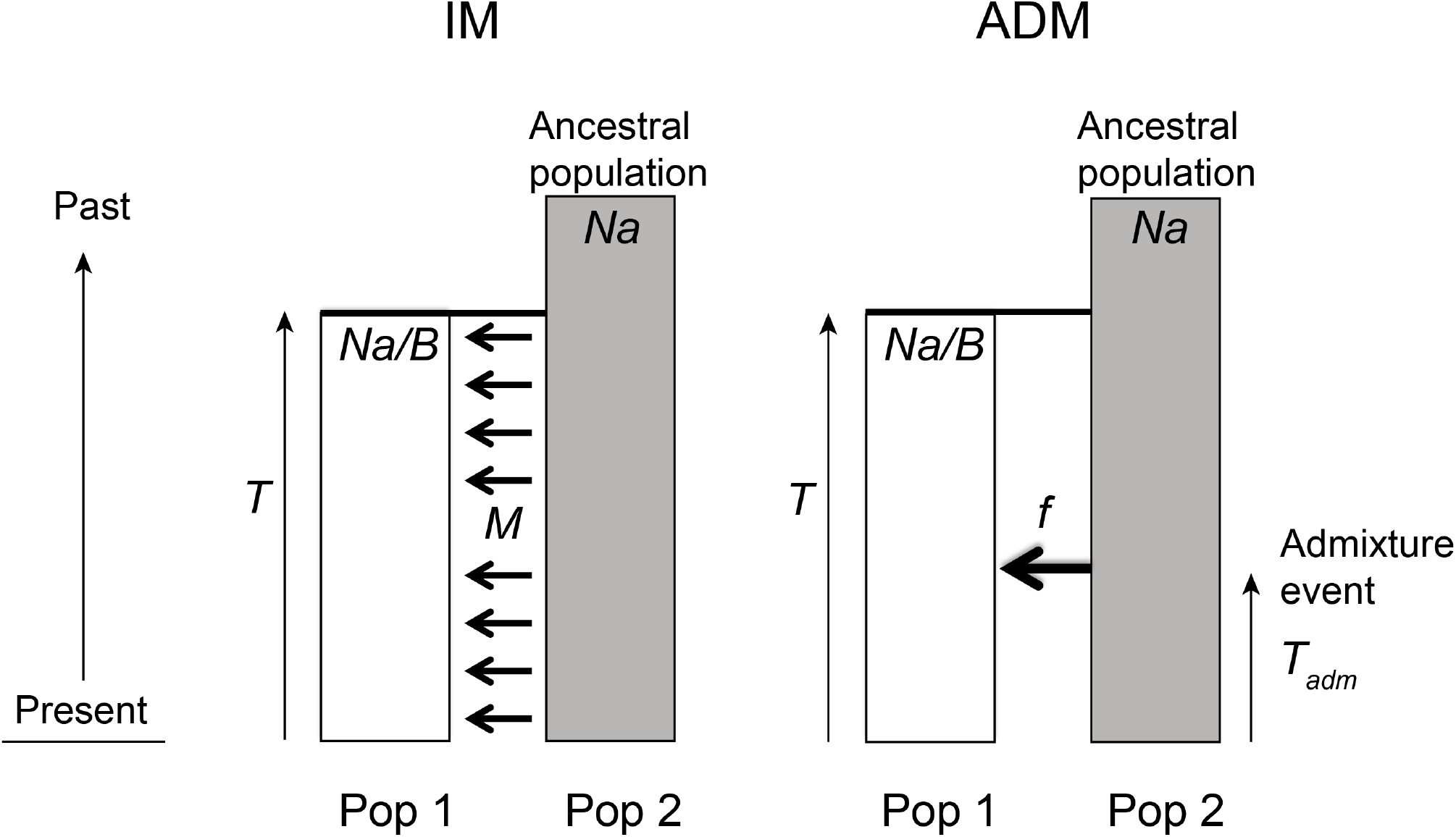
The IM (divergence with continuous migration) and ADM (divergence with instantaneous admixture) models of population divergence with gene flow, showing the demographic parameters estimated in our blockwise method analyses. Gene flow can be modelled in either direction*. Na* represents the size of an ancestral population extending back into the past that splits into two daughter populations at time *T* (scaled by 2*N_e_*) generations. One population retains the same population size *Na,* and one is free to have a new population size *Na*/*B,* where *B* is a scaling factor. Post divergence gene flow in the IM model is a continuous process with total *M* per generation = *4N_e_ * m,* where *m* is the individual migration rate per generation. In the ADM model, gene flow is modelled as an instantaneous admixture event at time *T_adm_* (scaled by 2*N_e_*) generations ago, at which a fraction *f* of lineages in the source population are transferred to the receiving population.

Discriminating among alternative models of gene flow and estimating the relevant demographic parameters are data-hungry problems and a growing number of approaches exploit the signal contained in whole genome data (WGD) for a small sample of individuals (Li & Durbin, 2011; Lohse & Frantz 2014; Bunnefeld et al., 2018). Whilst these approaches are currently applicable to a limited diversity and complexity of demographic models (discussed further below), their requirement for only a small number of relatively low quality genomes makes them accessible and affordable for non-model taxa, including rare taxa for which larger samples of individuals and reference genomes often do not exist (Allendorf et al., 2010; Fuentes-Pardo & Ruzzante 2017).

Here we use two such approaches to infer the population divergence history of *Pleistodontes nigriventris*, a wasp that is the only pollinator of an endemic fig species (*Ficus watkinsiana*) found in two widely separated blocks of rainforest along the east coast of Australia (see below) (Dixon 2003; Lopez-Vaamonde et al., 2002). Our overall objective is to infer the extent and direction of fig wasp gene flow between these two populations, using WGD for just two individuals per population. Our approaches take advantage of the haplodiploidy of the Hymenoptera, by sampling males whose haploid genomes facilitate data analysis and interpretation.

We first compare support for alternative IM and ADM models using a parametric maximum-composite likelihood method (Lohse et al., 2011; 2016), and then compare these results with those obtained using the Pairwise Sequentially Markovian Coalescent (PSMC) (Li & Durbin, 2011), a non-parametric method. Both are well-suited to the pairwise population divergence hypotheses we explore in *Pleistodontes nigriventris*. We chose these methods because they infer population history based on different aspects of genome-wide sequence variation, and have contrasting limitations. The composite likelihood framework developed by Lohse et al. (2016) is based on a blockwise summary of sequence variation, while PSMC exploits the information contained in the density of pairwise differences along a minimal sample of two haploid genomes. The blockwise method – by design – lacks power to detect very gradual demographic changes (for example, in population size), while PSMC is known to smooth out very sudden changes (Li & Durbin, 2011). The two methods therefore complement each other and together provide a comprehensive picture of population history.

Fig trees are keystone biological resources in tropical and subtropical habitats worldwide (Cook & Rasplus 2003; Harrison 2005), and fig fruits and their insect inhabitants are important model systems in the study of community assembly and coevolution (Cook & Rasplus 2003; Segar et al., 2014). Our target species *Pleistodontes nigriventris* is the specialist pollinating wasp of *Ficus watkinsiana* (Lopez-Vaamonde et al., 2002; Male & Roberts 2005, Rønsted et al., 2008), a monoecious rainforest fig restricted to northern and southern populations over 1000km apart along the eastern coast of Australia (Fig. 2) (Dixon 2003). While the existence of intervening *F.* watkinsiana trees and associated *P. nigriventris* cannot be categorically excluded, no populations are known; the demographic history of *P. nigriventris* can thus be modelled in terms of pairwise population divergence.

**Figure 2.**
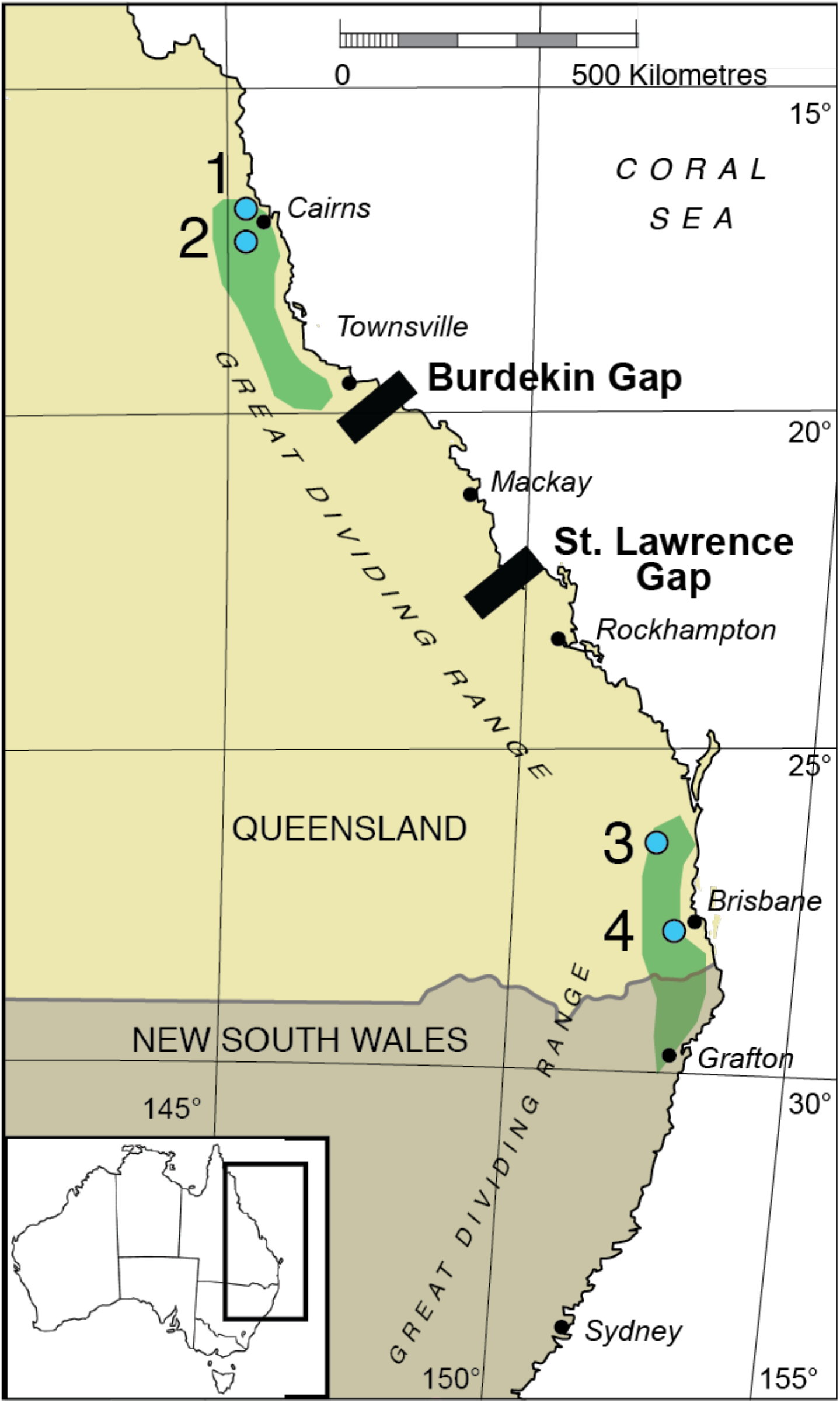
Map of the east coast of Australia, showing the distribution of Ficus watkinsiana (green) (after Dixon (2003)), the Burdekin and St. Lawrence Gaps (black), and the four sample sites (blue circles). The two Northern sampling sites are Kamerunga (1) and Kairi (2); the two Southern sites are Settlers Rise (3) and Main Range (4). The location of the study region within the whole of Australia is shown by the box in the inset at bottom left.

The distribution of Australian rainforests has been dictated by two major processes. First, falling temperatures during the Miocene restricted them to areas of higher rainfall along the eastern and southern coasts (Markgraf et al., 1995), separated from drier inland habitats by the Great Dividing Range (Chapple et al., 2011) (Fig. 2). Second, during the Pleistocene climate oscillations, east coast Australian rain forests repeatedly expanded from, and contracted into, a latitudinal series of refugia separated by intervening areas of dry forest and shrublands (Bryant & Krosch, 2016; Chapple et al., 2011). The rainforest areas occupied by *F. watkinsiana* are currently separated by two major dryland corridors (Fig. 2): the Burdekin Gap, located between Mackay and Townsville, is the largest dry land corridor on the east coast, and the St. Lawrence Gap is a smaller lowland dry corridor located 350km further south (Weber et al., 2014; Bryant & Krosch, 2016). While the formation and stability of these dryland corridors through time is incompletely characterised (Bryant & Krosch, 2016), both have been implicated in restricting dispersal and driving population divergence in rainforest plants (Burke et al., 2013), including *F. watkinsiana* (Dixon, 2003H; Haine & Cook, 2005), and animals (e.g. Schauble & Moritz, 2001; Pope et al., 2001; Nicholls & Austin, 2005; Brown et al., 2006; Dolman & Moritz, 2006; Baker et al., 2008, MacQueen et al., 2012; Rix & Harvey, 2012; Bryant & Fuller, 2014; Bryant & Krosch, 2016).

The impact of biogeographic barriers on the genetic makeup of any species depends on how long ago population divergence occurred, the sizes of the populations, and the direction, mode and frequency of gene flow between them (Aeschbacher et al., 2017; Ringbauer et al., 2018). Previous studies of fig/fig wasp systems provide evidence for two contrasting paradigms with which patterns in *Pleistodontes nigriventris* can be compared. Though female fig wasps are poor active flyers (and the males are wingless and do not leave their natal fig) (Ware & Compton 1994a,b), they can be dispersed over large distances by wind currents (Ahmed et al., 2009; Liu et al., 2015), particularly when emerging from fig fruits high in the forest canopy (Harrison & Rasplus, 2006; Kobmoo et al., 2010; Yang et al., 2015; Liu et al., 2015; Sutton et al., 2016). Long range dispersal by fig wasps is supported by lack of genetic structure over distances ranging from several hundred to >1500 km in pollinating fig wasps (Kobmoo et al., 2010; Liu et al., 2015; Tian et al. 2015; Bain et al., 2016) and in host figs - particularly monoecious species that are often large trees (Nazareno et al., 2013; Bain et al., 2016). *Ficus watkinsiana* can grow to 50m, and if *Pleistodontes nigriventris* benefits from wind-assisted dispersal, we might expect to find significant gene flow between *F. watkinsiana* populations. In contrast to this pattern, local adaptation of diverging populations to local host figs and/or abiotic conditions could drive genetic divergence between fig wasp populations over similar or even relatively small geographic scales. Genetic divergence has been documented over tens of km between mainland and island populations of a Chinese fig wasp (Tian et al., 2015), and between the same northern and southern rainforest habitats studied here for two *Pleistodontes* pollinators of another monoecious fig, *Ficus rubiginosa* (Darwell et al., 2014).

Here we answer the following questions for *Pleistodontes nigriventris*:

1. Is there evidence of genetic divergence between populations either side of the Burdekin and St. Lawrence Gaps?
2. Is there a signal of post-divergence gene flow, and if so, in which direction?
3. Can we discriminate between the IM and ADM models of gene flow?
4. Do the blockwise and PSMC methods infer concordant population histories?
5. Is the inferred divergence time for *P. nigriventris* concordant with estimates for other co-distributed taxa?

## Materials and Methods

### Sample Collection

Samples were collected between January 2001 and August 2009 from four sites in Queensland, two in each of the northern and southern ranges of *F. watkinsiana.* The two Northern individuals were sampled from Kairi (N1: 17.21° S, 145.55° E) and Kamerunga (N2: 16.87° S, 145.68° E) and the two South individuals were sampled from Settlers Rise (S1: 27.68° S, 153.26° E) and Main Range (S2: 28.07° S, 152.41° E) (Fig. 2). Near-to-ripe *F. watkinsiana* fruits were collected and placed individually into specimen pots. Our sampling targeted male fig wasps, whose haploid genome facilitates analysis (Bunnefeld et al., 2018; Hearn et al., 2014). Once the wasps started to emerge (12-24 hours after collection depending on fig ripeness) figs were dissected and live males were placed directly into 70% ethanol to preserve for DNA extraction. Due to potentially high levels of sib-mating in fig wasps (Greeff et al., 2009; Sutton et al., 2016) all individuals were sampled from different figs.

### DNA extraction and sequenced-based confirmation of identity

DNA was extracted from whole male wasps 1.5-1.6mm long (Lopez-Vaamonde et al., 2002) using the Qiagen DNeasy Blood and Tissue Extraction kit. The Purification of Total DNA from Animal Tissues (Spin-Column) Protocol was followed with the following modifications to maximise DNA yield from these extremely small wasps. Step 1: Individual wasps were placed in 180 μl of buffer ATL and crushed using a mini-pestle. Step 2: Riboshredder RNase (Epicentre) was used in place of RNase A. Steps 7-8: Buffer EB was used in place of Buffer AE for the elutions as Buffer AE contains EDTA, which will interfere with the downstream library preparation. Extractions were eluted in smaller volumes (25 μl) than recommended and samples were incubated for longer (5 minutes). This protocol yielded 35.7-58.3 ng of DNA per wasp, despite their very small size (total body length *ca.* 1mm).

Identification of male fig wasps was confirmed by comparison of sample sequences to voucher sequences for a 433 base pair (bp) fragment of the mitochondrial cytochrome b (cytb) gene (Lopez-Vaamonde et al., 2001). Sequences were amplified using the primers CB1/CB2 (Jermiin & Crozier, 1994). 0.3 μl of DNA extraction was used per PCR reaction. The remainder of the PCR mix consisted of 2 μl BSA (10 mg/ml), 2 μl 10X PCR buffer, 0.8 μl MgCl2 (50mM), 0.3 μl of each primer (20 μM), 0.16 μl dNTPs (each 25 mM) and 0.1 μl Taq (Bioline 5U/μl), made up to 20 μl with autoclaved MilliQ water. Amplification was carried out using a Bio-Rad S1000 thermal cycler for 2 minutes at 94°C, 35 cycles of 30 seconds at 94°C, 30 seconds at 48°C, 40 seconds at 72°C, and a final elongation step of 5 minutes at 72°C. PCR products were visualised on a 2% agarose gel and cleaned using a shrimp alkaline phosphatase and exonuclease 1 protocol. 2.5 μl of SAPExo1 mix (1.425 μl SAP dilution buffer, 1 μl SAP (1U), 0.075 μl Exo1 (1.5U)) was added to each sample before being incubated for 40 minutes at 37°C followed by 15 minutes at 94°C. Only the forward strand was sequenced for each individual, using BigDye chemistry on an ABI 3730 machine at the Edinburgh Genomics facility. Sequences for the four individuals have been deposited on GenBank (individual accession numbers: N1 MF597824; N2 MF597800; S1 MF597825; S2 MF597826).

### High-throughput library preparation and sequencing

We generated an Illumina Nextera genomic library for each individual male fig wasp following the manufacturers’ instructions. To make best use of paired end sequencing, the library fragment size distribution should be unimodal with a majority of fragments longer than the 150bp read length. We checked the fragment size distribution by running 1μl of each library on a high sensitivity DNA Bioanalyzer chip (Agilent 2100). Libraries for each individual were end-labelled using a unique pair of indices, pooled and sequenced in one lane of 150bp paired-end reads on the Illumina HiSeq platform at the Edinburgh Genomics facility. Pooling volumes were calculated to achieve ~6-fold coverage per individual assuming 50 gigabases of data per lane and a genome size of 300 Mb based on an estimate for another Agaonid fig wasp, *Ceratosolen solmsi* (278 Mb; Xiao et al., 2013). Our strategy in balancing sequencing depth across individuals was to maximise data availability for our analyses while minimising cost, and to avoid biases in variant calling that could result from sequencing one or more individuals to substantially greater depth. Final coverage for each individual matched expectations and is shown in Table S2. The short read data have been deposited at the ENA short read archive (Cooper et al., 2020a).

### Bioinformatic pipeline

Reads for all four individuals were screened to remove low quality and contaminant reads and combined. After exclusion of contaminants, reads for all four individuals were combined to generate a *de novo* meta-assembly for *P. nigriventris* using *SPAdes* (version 3.6.2) (Bankevich et al., 2012). Reads from each individual were mapped back to this reference using BWA; variants were called using *GATK* (Van der Auwera et al., 2013; version 3.5.0) haplotype caller. After masking repeat sequences, sites with a minimum coverage of two and a mapping quality (>20) and base quality (>10) were identified using the *GATK* tool *CallableLoci*. The bioinformatic pipeline is summarised diagrammatically in Supplementary information, Figure S1.

#### (i) Quality control and processing of sequencing reads

To exclude low quality sequence, reads were checked using *Fast-QC* (www.bioinformatics.babraham.ac.uk/projects/fastqc) and trimmed at a base quality score of 20 (sliding window 15:20 and min length 50). Adapters were trimmed using *Trimmomatic* version 0.32 (Bolgar et al., 2014). Any adaptors found to be present after a second pass through *Fast-QC* were removed using *Cutadapt* (Martin, 2011). Paired-end reads were merged (assembled into pairs) using *PEAR* version 0.9.0 (Zhang et al., 2014) for *de novo* assembly only.

#### (ii) Filtering of contaminant sequences

Genomic data obtained from whole organism libraries commonly contain reads from associated non-target organisms, including symbionts, parasites and commensals. *Wolbachia* bacteria are common endosymbionts in fig wasps (Haine and Cook, 2005) and diverse fungal taxa have been identified from a genome assembly of the fig wasp *Ceratosolen solmsi* (Niu et al., 2015). To remove contaminant sequences from our data, we first aligned reads across all 4 individuals to create a single *Velvet* reference assembly (version 1.2.10) with a k-mer length of 31 (Zerbino and Birney, 2008). The filtered reads for each individual were then mapped to this *Velvet* reference assembly using *Bowtie2* version 2.2.3 (Langmead and Salzberg, 2012). Contaminant sequences were identified using blobtools (Kumar et al., 2013), which uses BLAST searches to create Taxon-Annotated-GC-Coverage plots (Blobplots) that allocate aligned reads to taxa at a user-specified level. We used four BLAST approaches to assess taxonomic matches: (i) The fast protein aligner *Diamond* (version 0.7.9) (Buchfink et al., 2015) was used alongside three BLAST (version 2.2.29) searches: (ii) against the NCBI nucleotide database (https://www.ncbi.nlm.nih.gov/nucleotide/), (iii) against the genome of the Agaonid pollinating fig wasp, *Ceratosolen solmsi* (Xiao et al., 2013), and (iv) against the genome of the pteromalid parasitoid wasp *Nasonia vitripennis* (Werren et al., 2010). Non-Arthropod reads (primarily allocated to Proteobacteria and Ascomycota) were excluded from further analysis.

#### (iii) Generation of a P. nigriventris reference assembly

After exclusion of contaminants, reads for all four individuals were combined and re-assembled using *SPAdes* (version 3.6.2) (Bankevich et al., 2012). The quality of this reference assembly was assessed using *BUSCO* (*Benchmarking Universal Single-Copy Orthologs*) version 1.1b1 (Simão et al., 2015) using the Arthropoda *BUSCO* set (http://busco.ezlab.org).

#### (iv) Variant calling

Filtered, merged reads from each individual were mapped back to the reference meta-assembly using the *Burrows-Wheeler Aligner* (*BWA)* version 0.7.10 (Li & Durbin, 2009). FASTA file indexes and sequence dictionaries were created using *samtools* (Li et al., 2009) *faidx* (version 1.2) and the *picard* (https://broadinstitute.github.io/picard/index.html) tool *CreateSequenceDictionary* (version 1.141) respectively. The BAM files were sorted and merged to create a single species BAM file using the *picard* tool *MergeSamFiles*. Duplicate reads were removed from these merged files using the *picard* tool *MarkDuplicates*. Variants were called using *HaplotypeCaller* in *GATK*. As male wasps are haploid, ploidy was set to 1 and the ‘-emit variants only’ option was used. Only SNPs were considered.

#### (v) Masking repetitive regions

Regions of repetitive DNA can cause assembly and mapping errors in short read data (Treangen & Salzberg, 2011). We created a library of repetitive regions in the reference meta-assembly using *RepeatScout* (version 1.0.5) (Price et al., 2005), and masked these using *RepeatMasker* (version open-4.0.6) (Smit, AFA, Hubley, R & Green, P. *RepeatMasker Open-4.0*. 2013-2015 http://www.repeatmasker.org). *RepeatMasker* outputs an annotation file which was used to create a BED file of repeat positions for downstream processing.

#### (vi) VCF filtering

High quality sites were identified based on coverage and mapping quality using the *GATK* tool *CallableLoci* and default parameter values except for the following: minimum base quality of 10, minimum mapping quality of 20, minimum read depth of two reads per site. The base quality score recalibrated (BQSR) BAM file was subsampled to extract BAM files for each individual. We used *CallableLoci* to generate BED files of callable regions in each individual. Repeat regions were excluded and positions meeting filters across all four individuals extracted using *bedtools multiIntersectBed*. This BED file was used to filter the VCF files (generated from the *GATK* pipeline) for callable variable sites using *bcftools* (Li, 2011) version 1.2.

### Fitting of IM and ADM models

#### (i) Specification of alternative IM and ADM divergence models

Both the IM and the ADM model assume an ancestral population with a constant effective population size (*N_a_*) that splits into two populations (North and South). One of the descendant populations maintains the same ancestral population size *N_a_*, whilst the other is free to change to (*1/B)* x *N_a_* (i.e. *B* scales the rate of coalescence in the other population). In both models (Fig. 1), population divergence occurs at time *T* in the past. In the IM model divergence is followed by continuous unidirectional migration at rate *M = 4Nm* migrants per generation (m is the per lineage probability of migrating). In the ADM model (Fig. 1), gene flow occurs through an instantaneous and unidirectional admixture event at time *T_adm_* which transfers a fraction *f* of lineages from the donor population into the recipient (Fig. 1). All time parameters are scaled in 2N_a_ generations. We assessed support for all four possible combinations of population size and gene flow under both the IM and ADM models (Fig. S2), as well as for simpler nested models which either involving gene flow but a single population size parameter (B=1) or two population sizes but no post-divergence gene flow.

#### (i) Generation of blockwise site frequency spectra (bSFS)

We used the composite likelihood calculation described in Lohse et al. (2016) to fit the IM and ADM models (see also Jordan et al., 2017 and Nürnberger et al., 2017). The method uses information in patterns of linked sequence variation contained in short blocks and has previously been used for demographic inference under a variety of demographic scenarios including the IM model (Lohse et al., 2012) and models of discrete admixture (Bunnefeld et al., 2018). For a sample of four haploid individuals (two from each of the Northern and Southern populations), we can distinguish four mutation types: variants in the two Northern samples (Nvar), variants in the two Southern samples (Svar), fixed differences between North and South (Fixed differences), or variants shared between North and South (Sharedvar) (Lohse et al., 2016). The site frequency spectrum of a block (bSFS) consists of counts of these four site types, i.e. vector {Nvar, Svar, Sharedvar, Fixed}. We used the automated recursion implemented by Lohse et al. (2016) to obtain the generating function of genealogies under the ADM model and computed the composite likelihood of both the ADM and IM models in *Mathematica* as described in Lohse et al. (2016) and Nürnberger et al. (2017).

The likelihood calculation assumes an infinite sites mutation model and no within-block recombination. Given these assumptions, blocks that contain both fixed difference and shared variants (violating the 4-gamete test) are not possible and were excluded from the likelihood calculation. We chose a block length that strikes a balance between potential bias (arising from recombination within blocks) and power (which suffers when too few blocks contain multiple variant sites). We used a custom python script to extract aligned sequence blocks of a fixed length of 387 base pairs, which corresponds to an average of 1.5 variant sites per block (see Table S1). The blockwise data were analysed in *Mathematica* (Wolfram Research, Inc., Mathematica, Version 10.4, Champaign, IL (2016)) (Cooper et al., 2020b). The computational cost of calculating composite likelihoods increases with the number of unique bSFS configurations considered in the data. We limited the number of bSFS configurations by lumping mutation counts above a threshold k_max_= 2 for the Nvar, Svar and SharedVar and k_max_= 3 for fixed differences. Since 87% of the blockwise data are within these k_max_ bounds and so included in the composite likelihood calculation exactly, the expected loss of power is minimal.

#### (iii) Estimation of demographic parameters

We estimated all model parameters for both the IM and ADM model. Estimation of *N_a_ (as N_a_ = theta /(4 μ;))* and scaling of *T* parameters into years requires an estimate of the mutation rate per generation (*μ*) and the number of fig wasp generations (*g*) per year. In the absence of any fig wasp estimate of *μ;*, we used the per generation mutation rate of 2.8 x 10^−9^ for *Drosophila melanogaster* (Keightley et al., 2014). Note that the relative timing of events in different models is not affected by this calibration. We assumed four fig wasp generations per year (*g=4*) and converted time estimates into years by multiplying by 2*N_a_* and dividing by g. The generation time for *P. nigriventris* is not known with certainty, and we assume 4 (range of 2-6) generations per year based on the following rationale. Duration of any single fig wasp generation can be estimated by measuring the time from pollination to ripening of individual figs. This is because wasp eggs are laid at the time of pollination, the emerging daughters disperse at the same time as the fig ripens, and these new females lay eggs rapidly as they live only a day or two as adults (Sutton et al. 2018). Estimates of the length of fig tree reproductive events are available for several species (e.g. Bronstein 1990), but not available for *P. nigriventris*. Jia et al. (2008) estimated a generation time of about 40 days for another Australian *Pleistodontes* fig wasp (*P. imperialis*) pollinating *Ficus rubiginosa*. This might suggest 8-9 generations per year for *P. imperialis*, but while these monoecious figs (including *F. watkinsiana*) fruit asynchronously year-round (Harrison, 2005), they are in highly seasonal environments and generation time can be much longer in cooler or drier periods. Fruit development time is also longer in species producing larger figs. Fruit development takes 3-8 months in winter figs of *F. macrophylla* (JMC, unpublished data), which has smaller figs than *F. watkinsiana* (Al-beidh, 2010). Hence, a range of 2-6 generations per year seems likely for *P. nigriventris*. We discuss the consequences of uncertainty in our generation time estimates.

#### (iv) Correcting for the effects of linkage between blocks

The composite likelihood calculations do not account for linkage between blocks. Given the difficulty of assessing the level of linkage between blocks in highly fragmented assemblies, we incorporated linkage effects using a parametric bootstrap approach (Lohse et al., 2016). We used *msprime* (version 0.3.1 (IM) and 0.4.0 (ADM)) (Kelleher et al., 2016) to simulate 100 datasets assuming a recombination rate of 2.719 x 10^−10^ per base (estimated from the data, see below). Each simulated dataset had the same total length as the real data. To keep simulations computationally tractable, we partitioned each simulated genome into 20 pseudo-chromosomes of equal length. Because our simulations allow for recombination anywhere along a linear genome, this parametric bootstrap scheme allows us to quantify both the uncertainty in estimates due to linkage between blocks and the potential bias due to recombination within them. Fitting the inferred model to these simulated datasets, we obtained 95% confidence intervals for parameter estimates as ±1.96 standard deviations (of the analogous estimates on the simulated data).

#### (v) Model selection

We used an analogous parametric bootstrap scheme to determine whether the IM or ADM models provided a significantly better fit to our data. We fitted both models to each of 100 bootstrap datasets simulated under the best-fit IM model parameters. The distribution of differences in log composite likelihood (calculated as ΔlnCL = 2*(ADMlnCL-IMlnCL)) between the IM model and the ADM model represents an expectation of selecting the ADM when the IM is true (Type 1 error). We determined critical values from this distribution at a significance level of 0.05.

#### (vi) Recombination rate estimation

There are no estimates of either the per generation recombination rate (*r*) or the population recombination rate (*ρ* = 4*N_e_r*) for any fig wasp species. We therefore estimated a recombination rate for *P. nigriventris* using a two-locus generating function that co-estimates *ρ* (scaled by *2N_e_*) and *θ*, for a sample of two haploid genomes drawn from a single panmictic population (here, the two Northern individuals) (Lohse et al., 2011).

### PSMC analysis

We compared the results of our blockwise analysis with inference using the Pairwise Sequentially Markovian Coalescent (PSMC) (Li & Durbin, 2011). PSMC infers the effective population size history from a pair of genomes, which may either be sampled from a single diploid individual, or a pair of haploid individuals. PSMC provides a continuous estimate of demographic parameters through time. While PSMC does not allow direct inference of population divergence, comparison of the PSMC trajectories of two diverged populations (here, North and South) reveals when these shared the same population size and so were likely part of the same ancestral population.

PSMC uses the density of pairwise differences along the genome to infer a trajectory of population size through time. Input files were generated from the per-individual BAM files using *samtools mpileup* (version 0.1.19). The pileup file was converted to a VCF file in *bcftools* (version 0.1.19) and a consensus sequence (fastq file) per individual was generated using the *bcftools* utility *vcf2fq*. Consensus files contained 31,371 and 33,523 contigs for the Northern and Southern pairs respectively. The two consensus files per population were merged using the *seqtk* function *mergefa* (version 1.0) and converted to the PSMC input format using the PSMC function *fq2psmcfa* (version 0.6.5). Only contigs that contain >10kb of unfiltered bases, ≥ 80% of which pass a quality threshold score, here set to ≥20 were included (6,900 and 7,305 contigs for the North and South pairs respectively). We analysed each pair of individuals with PSMC (version 0.6.5) using the following parameter values: N = 30, t = 20, r = 10.3 and p = “4+60*1+4”. Each dataset was sub-sampled 100 times to generate bootstrap replicates using the PSMC utility *splitfa* (version 0.6.5). Results were calibrated using the same mutation rate and generation times given above.

## Results

### De novo genome assembly for *Pleistodontes nigriventris*

Nextera libraries were generated successfully for all four male *Pleistodontes nigriventris* (Figure S3). After initial filtering and trimming the joint assembly of reads for all four individuals in *Velvet* contained 173,180 contigs ≥200bp (Table S1). We identified contaminant reads from a range of non-Arthropod taxa (Figure S4). Individual assemblies contained between 5k and 149k reads identified as bacterial. For three individuals over 97% of these were attributed to *Wolbachia* (and 24% in the final individual) (Table S2). The relative fraction of *Wolbachia* reads was significantly higher in Northern compared to Southern individuals (Chi-squared = 95159, df = 1, p < 2.2e-16), consistent with previous findings of between-population differences in *Wolbachia* prevalence in *P. nigriventris* (Haine & Cook, 2005). Excluding contaminant reads resulted in between 11 and 22.5 million reads per individual (Table S1). Re-assembly using *SPAdes* (Cooper et al., 2020a) improved the contiguity (139,731 contigs ≥200bp, N50 of 9,643 bp) and the completeness of the final assembly: *CEGMA* scores: 93.15% complete and 99.19% partially complete, *BUSCO* scores for the reference Arthropod gene set: 74% complete, 4.5% duplicated, 16% fragmented and 8.4% missing. We note that the *P. nigriventris* assembly, although fragmented, has higher completeness (in terms of complete Core Eukaryote Genes) than the genome of *Ceratosolen solmsi* (*CEGMA*: 88% complete), the only published fig wasp genome (Xiao et al., 2013). Repetitive elements made up 21.4% of the reference assembly and were excluded from subsequent analyses.

Processing of the filtered and aligned reads (mean coverage per individual of 3.7x) identified 1,837,396 SNPs. The blockwise site frequency spectrum contained the four site types in the following proportions: Nvar 0.164, Svar 0.153, Fixed 0.673, and Sharedvar 0.0102. The average pairwise diversity per site (after filtering) for the North and South populations were very similar (π = 0.000884 and 0.000822 respectively). This is one of the lowest estimates of genetic diversity reported for any sexually reproducing arthropod (Leffler et al., 2012; Romiguier et al., 2014). Pairwise *Fst* was 0.74, indicating high differentiation between the Northern and Southern populations.

### Inferring divergence with continuous migration and admixture

Our block extraction protocol (Cooper et al., 2020b) resulted in 775,977 blocks of 387 base pairs. Of these, 0.8% (5,872) contained both shared variants and fixed difference, violating the 4-gamete test, and were excluded from blockwise analyses. *Isolation with continuous migration (IM).* Models incorporating post-divergence migration and different effective population sizes (*N_e_*) received substantially greater support than otherwise equivalent models with no migration and/or a single *N_e_* parameter (Table 1a). The best-supported direction of migration is from South to North, irrespective of whether we assumed a single *N_e_* (compare model IM3 to IM2) or two *N_e_* parameters (compare IM7+IM9 to models IM6+IM8). The scenario in which the South population retained the ancestral *N_e_* (IM9; Figure 3a) had highest support. Under this model, the split between the two populations occurred 177 (95% C.I. 172–182) thousand years ago (kya) and the *N_e_* for the ancestral/South population was estimated to be slightly higher (69k, 95% C.I. 65.4k-72.6k) than for the North population (58k 95% C.I. 54.3k-61.5k). We inferred a migration rate *M* from south to north of 0.071. Although low (1 migrant every 28 generations), this estimate was significantly greater than zero (95% C.I. 0.045 – 0.097).

**Table 1.** Maximum composite likelihood estimates (MCLE) of model support and demographic parameters under (a) IM and (b) ADM models. The best supported model is highlighted in bold with parameter 95% confidence intervals estimated by parametric bootstrap. n(*N_e_*) indicates the number of population size parameters in the model, and *N_a_* indicates the population(s) retaining the ancestral population size. Mig indicates the direction of gene flow in the model, with 0 indicating a model with no gene flow. Model support is measured relative to the best fit model in each class (IM9 in (a) and ADM10 in (b)). Likelihood ratio tests (LRT) were calculated as 2ΔlnL.

| Model | $n(N_e)$ | $N_a$ | Mig | $\Delta\ln L$ | LRT | $T$ | $B$ | $M$ |
| --- | --- | --- | --- | --- | --- | --- | --- | --- |
| IM1 | 1 | both | 0 | -22,877 | n/a | 3.938 | n/a | n/a |
| IM2 | 1 | both | N->S | -5,078 | 35,598 | 5.210 | n/a | 0.044 |
| IM3 | 1 | both | S->N | -1,122 | 43,509 | 5.533 | n/a | 0.057 |
| IM4 | 2 | N | 0 | -22,740 | 274 | 3.523 | 1.198 | n/a |
| IM5 | 2 | S | 0 | -21,298 | 3,157 | 3.808 | 1.055 | n/a |
| IM6 | 2 | N | N->S | -2,855 | 36,885 | 6.417 | 0.774 | 0.051 |
| IM7 | 2 | N | S->N | -500 | 44,479 | 6.193 | 0.872 | 0.061 |
| IM8 | 2 | S | N->S | -1,757 | 39,081 | 4.573 | 1.348 | 0.065 |
| **IM9** | **2** | **S** | **S->N** | **0** | **45,481** | **5.137 (4.784-5.489)** | **1.193 (1.089-1.296)** | **0.071 (0.045-0.097)** |
| IM9 sim |  |  |  |  |  | 0.299 | 5.150 | 1.197 |

(b) ADM (Instantaneous admixture) models
| Model | $n(N_e)$ | $N_a$ | Mig | $\Delta\ln L$ | LRT | $T_{adm}$ | $T$ | $B$ | $f$ |
| --- | --- | --- | --- | --- | --- | --- | --- | --- | --- |
| ADM1 | 1 | both | 0 | -6,349 | 37,230 | n/a | 4.969 | n/a | 0.015 |
| ADM2 | 1 | both | 0 | -6,349 | 37,230 | n/a | 4.969 | n/a | 0.015 |
| ADM3 | 1 | both | N->S | -4,995 | 2,634 | 2.194 | 6.122 | n/a | 0.296 |
| ADM4 | 1 | both | S->N | -1,110 | 10,478 | 1.970 | 6.095 | n/a | 0.257 |
| ADM5 | 2 | N | 0 | -6,536 | 36,583 | n/a | 4.193 | 1.313 | 0.014 |
| ADM6 | 2 | S | 0 | -5,632 | 35,508 | n/a | 4.631 | 1.152 | 0.017 |
| ADM7 | 2 | N | N->S | -1,645 | 9,784 | 1.642 | 5.195 | 1.329 | 0.257 |
| ADM8 | 2 | N | S->N | -172 | 10,919 | 2.003 | 6.872 | 0.857 | 0.256 |
| ADM9 | 2 | S | N->S | -1975 | 9,122 | 2.281 | 7.276 | 0.767 | 0.291 |
| **ADM10** | **2** | **S** | **S->N** | **0** | **11,263** | **1.656 (1.509-1.802)** | **5.643 (5.317-5.966)** | **1.181 (1.093-1.269)** | **0.239 (0.206-0.272)** |
| ADM10 sim |  |  |  |  |  | 0.302 | 1.653 | 5.604 | 1.184 |

**Figure 3.**
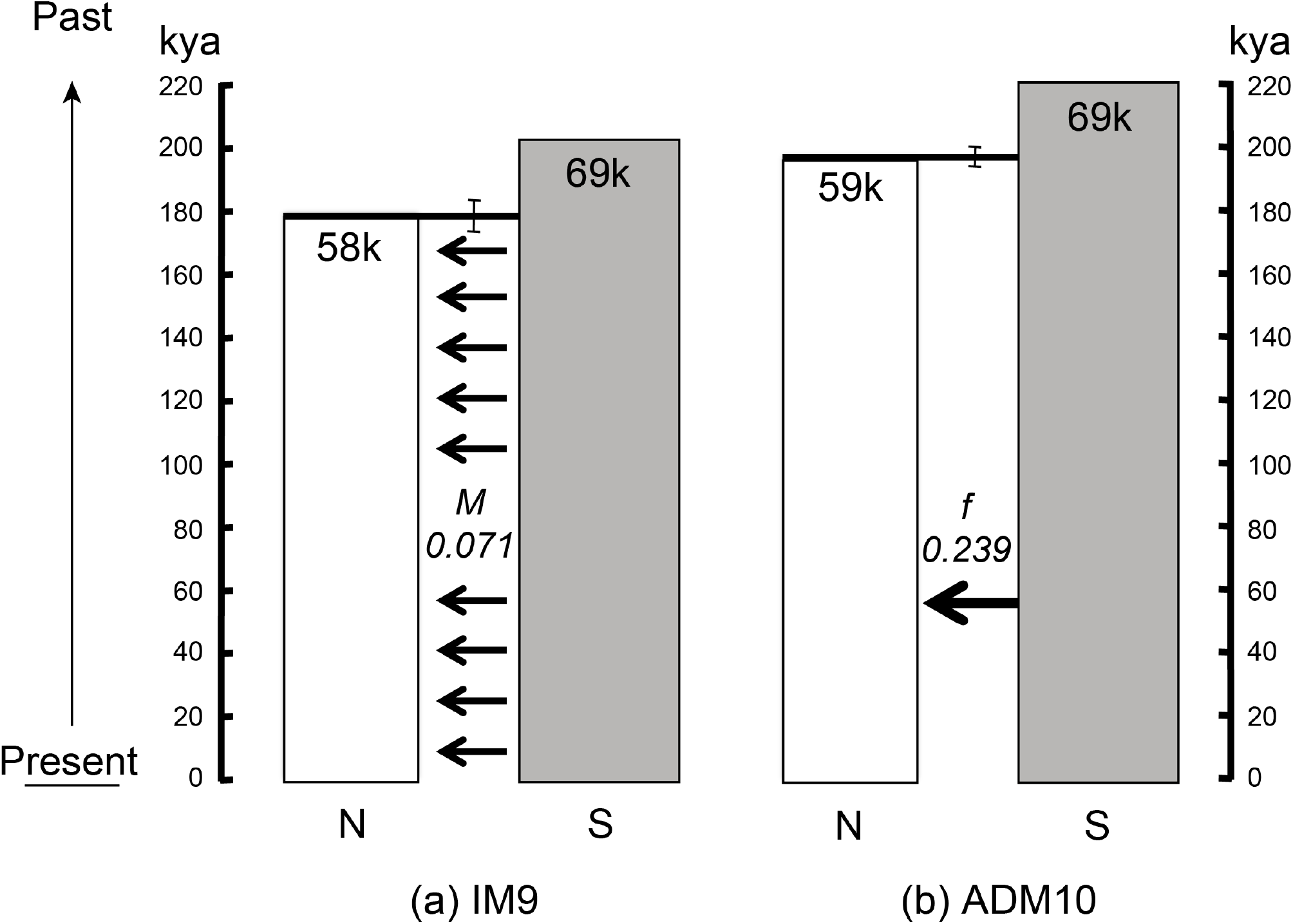
The best supported models under (a) IM and (b) ADM scenarios for *Pleistodontes nigriventris.* Bars joining the populations at divergence are the 95% confidence intervals for the divergence time. Time is measured in thousands of years, assuming 4 generations per year. The population widths are scaled according to their population size estimates (given in thousands). 95% confidence intervals for other parameters are given in Table 1.

#### Isolation with instantaneous admixture (ADM)

Inferences for models assuming instantaneous admixture mirrored those for IM models both in terms of model comparisons and parameter estimates: scenarios with gene flow and two *N_e_* parameters were significantly better supported than otherwise equivalent models without admixture and/or a single *N_e_* (Table 1b). The best-supported admixture direction was again from South to North, and the model assuming that the Southern population retained the ancestral *N_e_* (ADM10; Fig. 3a) had greatest support. *N_e_* estimates under the best fitting ADM history were similar to those under the analogous IM model (IM9): we inferred an ancestral/Southern *N_e_* of 69k (95% C.I. 66.1k-72.6k) and a lower Northern *N_e_* of 59k (55.2k-62.2k). In contrast, under the best fitting ADM9 the population split was estimated 196kya (95% C.I. 193-198kya), slightly older than under the analogous IM model. The admixture event was inferred to have occurred 57kya (95% C.I. 53-62kya) (Table 1b) and around a quarter (admixture fraction *f* = 0.239, 95% C.I. 0.206-0.272) of Northern lineages are inferred to have originated from the Southern population.

#### Greater support of instantaneous admixture

The ADM model had greater support than the IM model (ΔlnCL = 4175). Since the blockwise composite likelihood calculation ignores linkage between blocks and given that IM and ADM models are not nested, we cannot use a likelihood ratio test to compare the support for these models. To confirm whether the simpler IM model fits significantly worse than the ADM model we obtained a critical value for ΔlnCL (155.4 at p=0.05) using a fully parametric bootstrap. The difference in model support in favour of the ADM model ΔlnCL = 4175 in the real data far exceeds this and confirms that admixture provides a better fit to our data than continuous gene flow. To investigate which aspect of blockwise variation in the data allows discrimination between instantaneous admixture (ADM) and continuous migration (IM), we compared the frequencies of the most common bSFS configurations in the data with those expected under the best fitting IM and ADM models (Figure S6). Inspection of the residual reveals that the ADM model predicts both the frequency of monomorphic blocks ({0,0,0,0}) and blocks with more than three fixed differences ({0,0,>3,0}) better than the IM model.

### PSMC supports divergence and instantaneous admixture from South to North

We used PSMC (Li & Durbin 2011) to infer trajectories of *N_e_* change for Northern and Southern populations. Comparing the trajectories both with each other and with parameters inferred under the best fitting models of divergence and gene flow (ADM10) reveals a close correspondence between blockwise analyses and PSMC in several respects (Fig. 4). First, PSMC trajectories are consistent with the blockwise inference of a slightly larger *N_e_* in the Southern population compared to the Northern population. Second, the divergence time inferred by the blockwise analyses corresponds to a period in the PSMC at which the Northern and Southern populations show similar *N_e_* (overlapping confidence intervals), consistent with a shared ancestral population. Finally, PSMC infers an increase in the Northern *N_e_* prior to the time of admixture inferred under the ADM model. Such an increase in genetic diversity in the (Northern) recipient population is exactly what would be expected from a sudden admixture event. We note that PSMC also reveals an increase in the Southern *N_e_* around the same time which, however, is markedly smaller. While PSMC also shows an increase in *N_e_* in the very recent past (i.e. the last 3ky), the variation among bootstrap replicates (Fig. 4) suggests that our data lack the signal to reliably infer population size change over this timescale.

**Figure 4.**
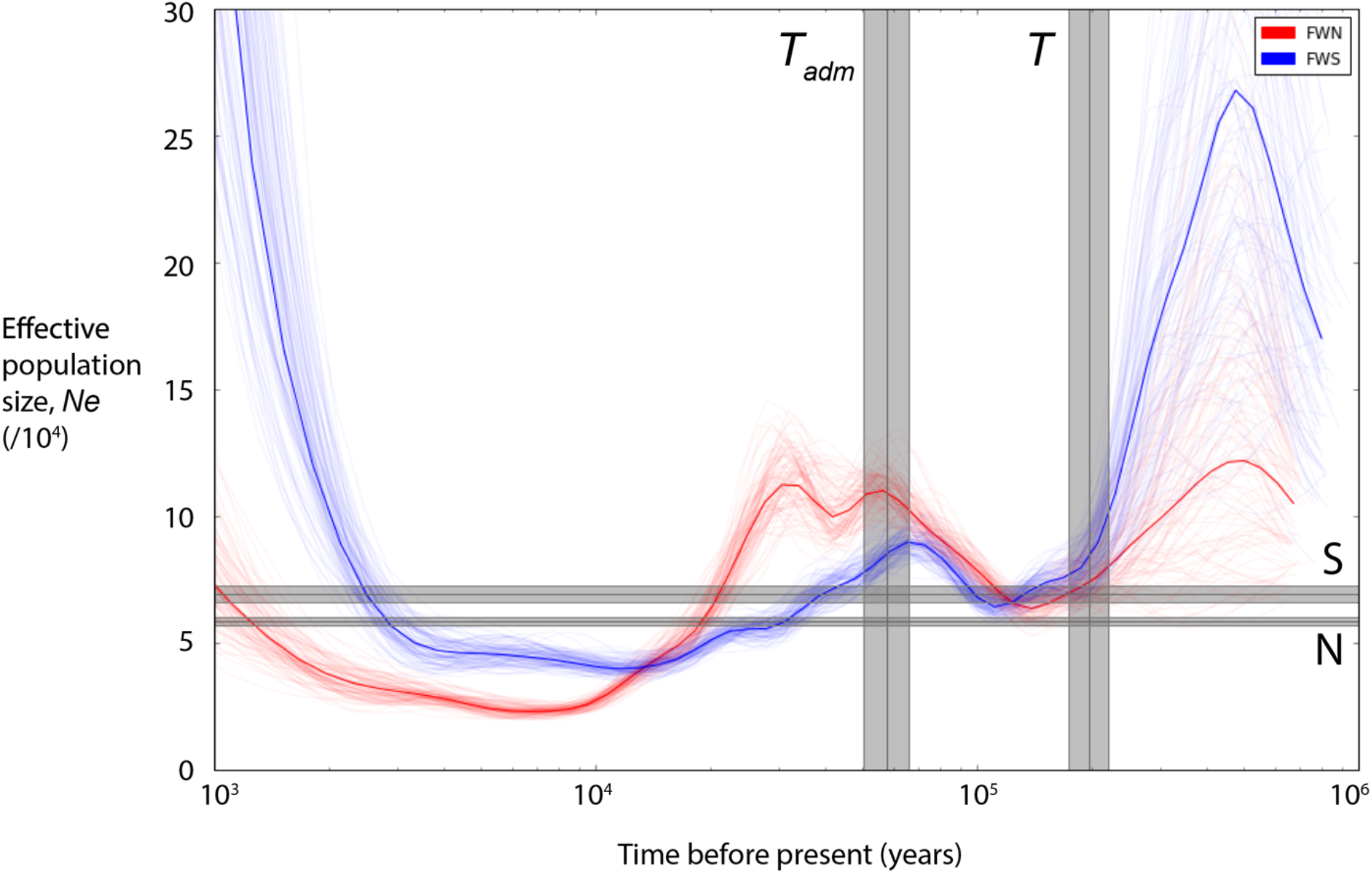
Population size trajectories for the Northern (red) and Southern (blue) populations inferred using PSMC. Heavy red and blue lines represent estimates for each population, thin lines show individual bootstrap replicates. Maximum composite likelihood estimate (MCLE) for the population sizes for the North and South populations and the dates of admixture (*T_adm_*) and population divergence (*T*) under the model that gives the best fit to the blockwise data (ADM10) (and their 95% confidence intervals) are overlaid in grey.

## Discussion

### Individual level population genomics of very small insects

Genomic data offer enormous potential in inference of population relationships and demography, but genomic resources remain limited for all but a tiny proportion of taxa. In some taxa, including rare species of conservation importance, it is also not possible to sample large panels of individuals (Allendorf et al., 2010; Fuentes-Pardo & Ruzzante 2017). In such cases, and where the distribution of populations is appropriate to a pairwise comparison, the approaches we use here allow demographic inference with minimal cost and sampling. Our approach may be of use in similar pair-wise population analyses of other east coast Australian forest taxa divided by the same dry habitat corridors, and other analogously distributed taxa.

We generated genomic libraries for individual male fig wasps, each only 1.5 mm long, resulting in over 770 thousand aligned blocks of sequence containing an average of 1.5 variable sites. Even with minimal samples of two haploid individuals per population, these whole genome data allowed estimation of demographic parameters with high confidence. A further advantage of the two methods we use is that neither makes any assumptions about phase in sequence data. Both can thus be applied to diploid organisms, provided that coverage is high enough to distinguish sequencing error from heterozygosity (ability to do so with high confidence is a major benefit of working with haploid organisms, including male Hymenoptera).

The best supported IM and ADM models gave similar estimates for the age of the initial divergence between Northern and Southern *P. nigriventris* populations, and their sizes. Further, we show that a burst of admixture (ADM) provides a significantly better fit than ongoing gene flow to the blockwise data and the individual population *N_e_* trajectories inferred by PSMC analysis. We first place our results in the broader context of phylogeographic work on taxa spanning the Burdekin and St. Lawrence Gaps, and then discuss the potential limits of our demographic inferences.

### Pleistocene population divergence across the Burdekin and St Lawrence Gaps in *P. nigriventris*

Australia has a complex climate history, a major feature of which is the aridification that started in the Miocene and continued throughout the Pliocene and Pleistocene (Schauble & Moritz, 2001; Martin, 2006; MacQueen et al., 2010, Frankham et al., 2016). This resulted in the gradual restriction of rain forests to the Eastern coast of Australia, separated from more arid inland habitats by the Great Dividing Range (Kershaw 1994; McGuigan et al., 1998; Schneider et al., 1998; Pope et al., 2000, Bell et al., 2007). Between 280 and 205kya, decreased precipitation and more severe aridification were associated with major faunal turnover in rainforest taxa (Hocknull et al., 2007). Against the backdrop of this general trend, the Pleistocene climate oscillations drove cycles of rain forest expansion during warmer, wetter interglacials and contraction during cooler, drier glacials (Byrne 2008; Maldonado et al., 2012; Burke et al., 2013). The current dry habitat corridors of the Burdekin and St. Lawrence Gaps are thought to be products of the long term aridification of Australia, and though it is uncertain when they first formed, they most likely existed through multiple Pleistocene cycles (Bryant & Krosch, 2016).

We found a strong signature of population divergence over the combined Burdekin and St. Lawrence gaps in *Pleistodontes nigriventris*. This is perhaps to be expected, given the obligate dependence of *P. nigriventris* on *Ficus watkinsiana,* and the restriction of this fig species to rain forest (Dixon 2003). However, our analyses do not allow inference of the ancestral distribution of *P. nigriventris,* and are compatible with either population being founded from the other, or vicariance of a previously continuous rainforest distribution (Martin, 2006). Assuming four generations per year for *P. nigriventris* (see below), the best fitting IM and ADM models for *P. nigriventris* both infer divergence between the Northern and Southern populations 170-200kya ago, in the Late Pleistocene.

In a review encompassing a wide range of plant and animal taxa, Bryant and Krosch (2016) identified a signature of population subdivision for the Burdekin Gap in 18 of 27 studies (nine of which have divergence time estimates) and for the St. Lawrence Gap in 10 of 23 studies (six of which have divergence time estimates). Pleistocene divergence across the Burdekin Gap has also been inferred for *Melomys cervinipes* (a wet forest rodent (Bryant & Fuller (2014)), *Petaurus australis* (yellow-bellied glider (Brown et al., 2006)), and *Varanus varius* (a large lizard with broad habitat preferences (Smissen et al., (2013)). Very few studies have considered insect population structure across the same potential barriers to gene flow. In contrast to our results for *P. nigriventris,* Schiffer et al., (2007) found no evidence for genetic divergence across the Burdekin gap in *Drosophila birchii*, a specialist rainforest fruit fly, but instead inferred a moderate gene flow across the whole range following a recent range expansion. Divergence estimates for other taxa spanning the Burdekin and/or St Lawrence Gaps are concentrated in the late Miocene to late Pleistocene (Bryant & Krosch 2016) with substantial variation across taxa. For example, divergence across the Burdekin Gap was estimated to have occurred in the early Miocene-late Oligocene >20 million years ago (mya) in *Uperoleia* frogs (Catullo & Keogh 2014; Catullo et al., 2014), and 31-51mya in assassin spiders (Rix & Harvey 2012). Given that most of these previous estimates are not based on any statistical model of population divergence, but rather a single (often mitochondrial) gene tree, it is unclear how much of the variation in previous divergence time estimates across these co-distributed taxa simply reflects coalescence variance and/or differences in calibration. Model based comparative studies are needed to assess whether past vicariance events have been shared in time across taxa with disjunct distributions across the Burdekin and St Lawrence gaps.

Two compatible explanations could explain observed population divergence in *P. nigriventris.* A first is that aridification and lack of suitable fig hosts in intervening habitats prevented viable dispersal between Northern and Southern populations. A second is that despite dispersal, migrants failed to contribute genes to receiving populations. This could happen if, following initial divergence, each of the Northern and Southern populations became locally adapted, but reciprocally maladapted (Rodriguez et al., 2017). Such failure could result from reciprocal mismatches in climate, or in coevolved interactions with co-distributed *Ficus watkinsiana.* That failure to disperse may not wholly explain divergence between Northern and Southern populations of *P. nigriventris* is suggested by inferred high dispersal within each population (low divergence between individuals), and the fact that some other fig wasps (including another *Pleistodontes* species (Sutton et al. 2016)) show little or no genetic divergence over distances of around 1000km, close to the separation between our populations (e.g. Kobmoo et al., 2010; Liu et al., 2015; Tian et al. 2015; Bain et al., 2016). Further evidence comes from lack of spatial genetic structure (and hence long range pollen and/or fruit dispersal) in other monoecious figs (e.g. Nazareno et al., 2013; Bain et al., 2016). That effects other than barriers to dispersal *per se* can structure intraspecific genetic diversity is also suggested by (potentially host-mediated) differentiation in other fig wasp species over distances of less than 50km (Tian et al., 2015). The extent to which Northern and Southern populations of *P. nigriventris* are able to interbreed and to induce galls in allochthonous host figs remains unknown, but could be tested experimentally.

### Effective population size of *P. nigriventris*

We estimated the effective population sizes for Northern and Southern populations of *P. nigriventris* to be ca. 60k and 70k respectively, with 95% confidence intervals over the IM and ADM models spanning 54k-73k. These figures fall within the range observed in other similarly sized chalcidoid and cynipoid Hymenoptera (e.g. Bunnefeld et al., 2018; Walton et al., 2020). The true census population size for *P. nigriventris* is almost certainly much higher, given that a single large fig tree can bear many thousands of pollinated figs, each of which required entry by at least one female *P. nigriventris*. Census population size (*N*, which contributes to fruit set) is generally one to several orders of magnitude greater than *N_e_* (Lewontin 1974; Frankham 1995; Palstra and Ruzzante 2008), particularly in taxa that, like fig wasps, show high levels of sib-mating (Kimura and Crow 1963; Leffler et al., 2012; Sutton et al., 2016 for *Pleistodontes*; Molbo et al., 2004 for *Pegoscapus*) and can experience large population fluctuations and genetic bottlenecks (Bronstein and Hossaert-McKey 1996; Harrison 2000; Wachi et al., 2016). Both are likely explanations for the similar and very low genetic diversities observed in each of the Northern and Southern populations (π = 0.000884 and 0.000822 respectively), amongst the lowest estimates for any sexually reproducing arthropod (Leffler et al., 2012; Romiguier et al., 2014).

### The Northern population of *P. nigriventris* received a burst of admixture from the South at the end of the Pleistocene

Both the best fitting ADM and IM models inferred gene flow from the Southern to the Northern population. While the continuous migration rate inferred under the best IM model (*M* = 0.071) is much lower than the estimated admixture fraction (f=0.24) under the better supported ADM model, the overall amount of gene flow inferred under both models is in fact very similar: Under the IM model, the probability that a single lineage sampled from the North is derived from the South via gene flow is 1-e^(-*MT*). Given our estimates for these parameters in model IM9, this is ~0.3, so very comparable to the admixture fraction inferred under the best ADM model. While both models agree in the overall amount of post-divergence gene flow, our model comparison clearly shows that genetic exchange occurred as a sudden burst rather than a continuous process. Additional support for a discrete admixture event comes from the contemporary increase in the Northern population size revealed by PSMC. The inferred combination of divergence and admixture is compatible with the following demographic scenario: (i) Northern and Southern populations were separated 170-200 kya following contraction of suitable habitat and expansion of intervening inhospitable dry forest corridors. (ii) Northern and Southern populations remained separated for over 100ky until favourable conditions in the late Pleistocene allowed expansion of one or both populations of *Ficus watkinsiana,* to the point at which substantial genetic exchange was possible between populations of pollinating *P. nigriventris*. (iii) Subsequent aridification resulted again in range contractions and a shutdown of gene flow.

Notwithstanding the uncertainty in our time calibration for *P. nigriventris* (see below), the date of the inferred admixture event falls in Marine Isotope Stage 3 (27-60kya), a period in the late Pleistocene characterised globally by abrupt phases of warming and cooling (Siddall et al., 2008; Van Meerbeeck et al., 2009). These warming phases were interspersed by cooler periods around every 7,000 years (Clark et al., 2007), and even during the cooler periods average temperatures were much higher than during the Last Glacial Maximum (~19-21kya) (Van Meerbeeck et al., 2009). The lower bound (53kya) of the inferred admixture time corresponds approximately to the warmest point in one of these cycles, and it is tempting to suggest that this allowed temporary expansion of the range of the *F. watkinsiana*/*P. nigriventris* mutualism and secondary genetic contact. Such climatic instability also raises the possibility that any selection in favour of locally adapted fig wasps could sometimes have been replaced by selection in favour of adaptive introgression of migrant genes (Hedrick 2013).

Our results do not rule out bidirectional dispersal, but rather imply a signal of predominant gene flow from the South into the North. This could indicate conditions that facilitated northwards dispersal in this direction, such as prevailing winds from the south. Records from the 1940s-2000s do indicate stronger winds from the south in eastern Australia (Australian Government, Bureau of Meteorology, http://www.bom.gov.au/climate/). However, it is not known whether current trends can be extended back into the past. Similar post-divergence dispersal from south to north across the Burdekin gap has been inferred in a small number of other rain forest-associated taxa (Bryant & Fuller, 2014; Bryant & Krosch 2016). Alternatively, the asymmetry in genetic exchange between North and South we infer may be the result of genetic incompatibilities that arose and became fixed during periods of isolation.

### Limits of demographic inference

Our inference of the population history of *P. nigriventris* is contingent on the realism of our models, ability to incorporate effects of linkage, and scaling of time estimates. We consider these issues in turn.

Our IM and ADM models assume two populations with a maximum of two different and constant *N_e_* parameters (Fig. 1). While this simple model is unlikely to be true for any organism, the assumption of two panmictic populations is, as far as is known, a good approximation for the current distribution of *P. nigriventris.* Fitting explicit models necessarily involves simplifying assumptions, and alternatives for limiting the number of N_e_ parameters (such as assuming equal N_e_ for all populations, or an even split between daughter populations) are even less realistic. Concordance in *N_e_* estimates between our IM/ADM models and our PSMC analysis (in which *N_e_* is free to vary through time) suggests that drastic changes in *N_e_* are not a feature of the population history of *P. nigriventris*, and that this simplifying assumption is unlikely to affect our inference. Agreement in parameter estimates across models and the close fit of our best ADM model to the observed blockwise data both suggest that we have captured key aspects of the demographic history of *P. nigriventris.* Perhaps the main result of our analyses is that we have shown that continuous and discrete gene flow can be clearly distinguished in this species, even over relatively recent timescales (both in terms of sequence divergence and genetic drift). While most analyses of demographic history choose *a priori* to model gene flow as either a continuous process or a discrete event, these two extremes of model space are rarely compared directly and it remains an open question which one better captures the demographic history in most taxa. All three demographic scenarios we have considered (IM, ADM and PSMC) are nevertheless crude simplifications of what is likely a more complex history, possibly involving repeated cycles of rain forest expansion and contraction. The upland wet forests south of the Burdekin Gap in the Clarke Range and Conway Peninsula are thought to have persisted through multiple Pleistocene climate cycles (Stuart-Fox et al., 2001), and it would be interesting in future to use the information contained in larger samples to explore more realistic histories involving repeated admixture pulses (Jésus et al., 2006) in *P. nigriventris* and *F. watkinsiana*, potentially involving additional intermediate populations that may no longer exist (e.g. Stone et al., 2017). Likewise, it would be interesting to test how well the blockwise distributions of divergence and diversity in *P. nigriventris* fit a more complex generalised class of IM models (GIM) that involve periods of historic gene flow (Costa & Wilkinson-Herbots 2017). Exploring these models would allow bridging of the model space between the IM and ADM histories we consider here (Costa & Wilkinson-Herbots 2017). However, unlike the bSFS framework we have used here, analytic results for GIM models are currently limited to pairwise samples, which, all else being equal, are less informative about gene flow (Lohse et al., 2016, Fig. 7).

Demographic analyses that are based on genome-wide samples of either single variants or loci/blocks generally assume that blocks are statistically independent, i.e. unlinked. A common way of dealing with linkage effects is to use subsampling, either by sampling a fixed minimum distance apart or by resampling bootstrap. We have instead opted for a fully parametric bootstrap that incorporates recombination within and linkage between blocks explicitly. Although this is computationally more intensive than subsampling, we believe it is the only way to accurately capture the effect on parameter estimates. Our parametric bootstrap for the best fitting IM and ADM models gave narrow 95% confidence intervals for all model parameters. Importantly, mean parameter estimates obtained from the simulation replicates are very close to the MLEs used to simulate the datasets (Table 1). This confirms that biases in parameter estimates due to violations of the assumption of no recombination within blocks are negligible, as shown in previous studies on a range of organisms (Jennings & Edwards, 2005; Lanier & Knowles, 2012; Hearn et al., 2014; Bunnefeld et al., 2015; Wang & Liu, 2016). Another standard assumption of demographic inference is that sequences evolve neutrally. While our approach sampled blockwise sequence variation genome-wide, one would expect both genome assembly and read mapping to be easier in regions under selective constraint. As a consequence, our blockwise dataset is likely enriched for conserved coding regions. In a methodologically similar study of a European gall wasp, Hearn et al., (2014) tested for the effect of selective constraint by partitioning blocks according to the proportion of coding sequence they contained, and scaling the estimated genome-wide mutation rate between values for synonymous and non-synonymous mutations. They found that mutation rate heterogeneity had no impact on the inference, reporting only a slight increase in *N_e_* and divergence time estimates. Given these results and the lack of annotated genomes or transcriptome data to aid gene detection, we have not pursued this here. However, it will be interesting to check whether selective constraint can explain the observed higher frequency of invariant blocks in our *P. nigriventris* data compared to expectations under both the best fitting IM and ADM models. Taken together, these results suggest that recombination within blocks and background selection had a minimal impact on our inference of demographic history.

Scaling divergence and admixture times into years requires knowledge of both the mutation rate and generation time. Both are uncertain for *P. nigriventris.* In the absence of direct mutation rate estimates for fig wasps, we used an estimate from *Drosophila melanogaster* (Keightley et al., 2014). While it is unclear how well this matches mutation rates in fig wasps, it is reassuring that the few published mutation rate estimates for other insects are similar (e.g. 2.9 x 10^−9^ for the butterfly *Heliconius melpomene*). A likely greater source of calibration uncertainty is average wasp generation time, which is not known for *P. nigriventris*. Our estimate of 4 generations per year is based on data for other Australian *Pleistodontes* species, taking into account the seasonal habitat and large (slowly developing) fruits of *F. watkinsiana* (see Methods). Calibration of the timing of demographic events in *P. nigriventris* scales simply and inversely with generation time, i.e., halving the number of generations per year doubles the age of events. Assuming 2 and 6 generations per year (rather than 4 as we have done here) still places divergence and admixture times for *P. nigriventris* in the late Pleistocene: for 2 generations/year, *T*=392(386-396)ky and *T_adm_*=114(106-184)ky, and for 6 generations/year, *T*=131(129-132)ky and *T_adm_*= 38(35-61)ky. However, in the absence of precise information our attempt to match the demographic history of *P. nigriventris* to past climatic events remains speculative.

Our results show that robust demographic inferences can be made for very small sample sizes of non-model taxa without significant associated genomic resources. While our motivation for assembling a genome for *P. nigriventris* was solely to generate a dataset for demographic inference, genome assemblies are arguably a more generally useful resource than other types of genomic data (e.g. RadSeq) and can be improved in the future (e.g. by adding RNASeq and long read data to improve annotation and contiguity respectively). Inference based on <u>*de novo*</u> assemblies and minimal sampling is likely to become the norm for taxa that are rare or hard to sample. Figs support highly structured (and often co-evolved) communities of pollinators, inquilines and natural enemies (Lopez-Vaamonde et al., 2001; Segar et al., 2014), and while a growing body of work addresses phylogeographic relationships in figs and their pollinators (Bain et al., 2016; Rodriguez et al., 2017; Yu et al., 2019), much less is known about the phylogeography of non-pollinating fig wasps (Sutton et al., 2016). Our approach provides a framework for comparative analyses that reconstruct the assembly of these species-rich communities. More broadly, given that many questions about both intraspecific demography history and speciation come down to distinguishing between ongoing gene flow and discrete admixture pulses, systematic power analyses on how this can best be achieved - especially for genomic data from non-model organisms without a contiguous reference genome - are urgently needed.

## Acknowledgements

This work was funded by a UK NERC grant (NE/J010499) to GS, KL and JMC. LC was supported by a PhD studentship from the UK NERC. KL was supported by a NERC fellowship (NE/L011522/1)

## Data Accessibility

The short read data and the *SPAdes* genome assembly have been deposited at the ENA short read archive (number PRJEB35527).

CO1 barcode sequences for the four *Pleistodontes nigriceps* individuals have been deposited in Genbank: N1 MF597824; N2 MF597800; S1 MF597825; S2 MF597826).

A *Mathematica* notebook and blockwise sequence data for this study are available from the Dryad repository, doi: (added on acceptance).

## Author contributions

LC, GNS, KL and JMC designed the research. LC, LB, KL and JH performed the research and analysed the data. GNS, KL and LC wrote the paper, with editorial input from all authors.

## Supplemental Information

**Table S1.** Summary statistics for the *Velvet* and *SPAdes* assemblies for the four individual *Pleistodontes nigriventris* fig wasps. Contig N50 is a measure of assembly quality, and indicates that 50% of bases in the assembly are contained in contigs greater than or equal to this value. The CEGMA ‘Complete’ % completeness indicates the percentage of the 248 core eukaryotic genes (CEGs) for which >70% of the protein length was present in the genome assembly the Core Eukaryotic Genes Mapping Approach (CEGMA) program (version 2.5) (Parra et al., 2007; Parra et al., 2009). The CEGMA ‘Partial’ % completeness indicates the summed percentage of CEGs scored as complete or represented by <70% of their protein length in the genome assembly. BUSCO scores represent the percentages of complete (C), duplicated (D), fragmented (F) and missing (M) BUSCOs (Benchmarking sets of Universal Single-Copy Orthologs) for the Arthropoda set of 2675 genes found in the assembly. Percentage masked is the percentage of each assembly identified as comprising repetitive elements. SNPs is the number of SNPs called during the GATK variant calling pipeline, excluding multivariate sites. π is the average pairwise nucleotide diversity per site across individuals.

| Statistic | Velvet | SPAdes |
| --- | --- | --- |
| No. of individuals | 4 | 4 |
| No. of contigs ≥ 200bp | 173,180 | 139,731 |
| Contig N50 | 4,104 | 9,643 |
| Mean contig length | 2,282 | 3,053 |
| Maximum contig length | 90,040 | 99,701 |
| Contig span (total length of contigs ≥ 200bp) | 395,224,362 | 426,733,192 |
| No. of scaffolds ≥ 200bp | | 105,765 |
| Scaffold N50 |  | 35,331 |
| Mean scaffold length |  | 4,097 |
| Maximum scaffold length |  | 581,070 |
| Scaffold span (total length of scaffolds ≥ 200bp) | | 433,370,639 |
| GC content | 0.293 | 0.296 |
| CEGMA - complete %completeness | 96.77 | 93.15 |
| CEGMA - partial %completeness | 99.19 | 99.19 |
| BUSCO C |  | 74.0 |
| BUSCO D |  | 4.5 |
| BUSCO F |  | 16.0 |
| BUSCO M |  | 8.4 |
| Percentage masked |  | 21.39 |
| SNPs |  | 1,837,396 |
| Mean nucleotide diversity across individuals ($\pi$) | | 0.00399 |
| Mean coverage for the species-wide reference assembly |  | 14.805568 |
| Mean coverage per individual |  | 3.701 |

---

**Table S2.** Summary statistics for mapping of paired reads for each individual to the SPAdes reference assembly. A phred-scaled mapping quality score of 20 (Q20) means the aligner has designated the read as having a 1 in 100 chance of being misaligned. Average read coverages per individual were calculated using Mosdepth (version 0.2.9) (Pedersen and Quinlan, 2018) and per-individual bam files. Average SNP coverages per individual were extracted using vcftools (version 0.1.16) and mean depth per individual calculated in R.

| Individual | Contaminant reads (% attributed to *Wolbachia*) | Total reads (including both of the pair) | Average read coverage per individual | Average SNP read depth per individual | Percentage of reads aligned (including both of the pair) | Reads that mapped with Q20 or higher as percentage of all aligned reads |
| --- | --- | --- | --- | --- | --- | --- |
| North 1 | 103,059 (99.75) | 17,947,652 | 6.32 | 6.46 | 99.98 | 99.73 |
| North 2 | 148,885 (97.80) | 22,452,403 | 7.17 | 7.64 | 99.97 | 99.71 |
| South 1 | 14,021 (23.90) | 10,994,084 | 3.75 | 3.89 | 99.93 | 99.66 |
| South 2 | 5,372 (99.20) | 16,653,787 | 5.32 | 5.85 | 99.97 | 99.74 |

## Supplementary Figures

**Figure S1.**
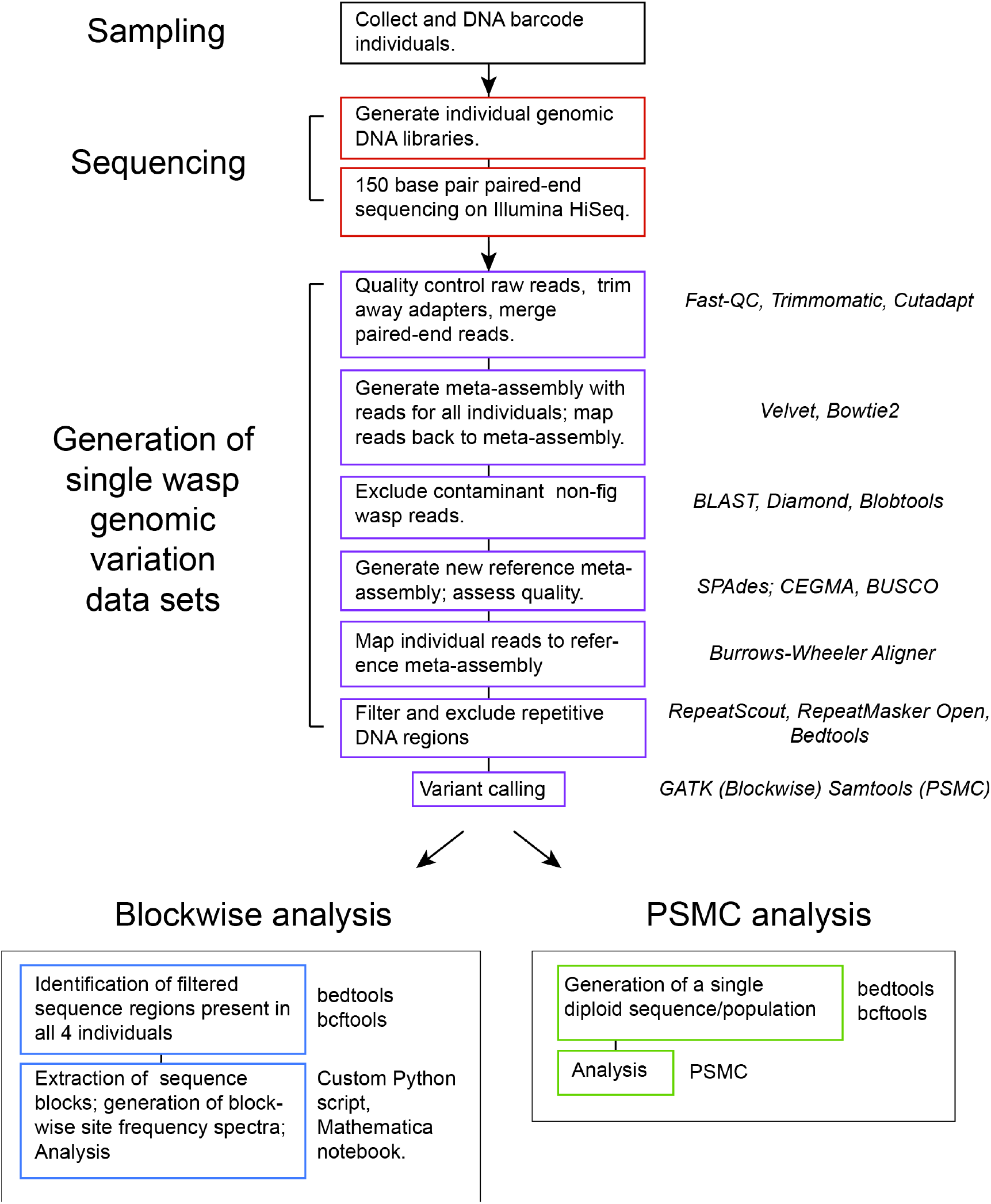
Diagrammatic summary of the informatic pipeline leading to the blockwise and PSMC analyses used in this study. Where a particular approach was used for one but not both analyses, the relevant analysis is specified in brackets. We encourage readers to update our pipeline in a modular fashion as newer (and superior) tools are developed for individual steps.

**Figure S2.**
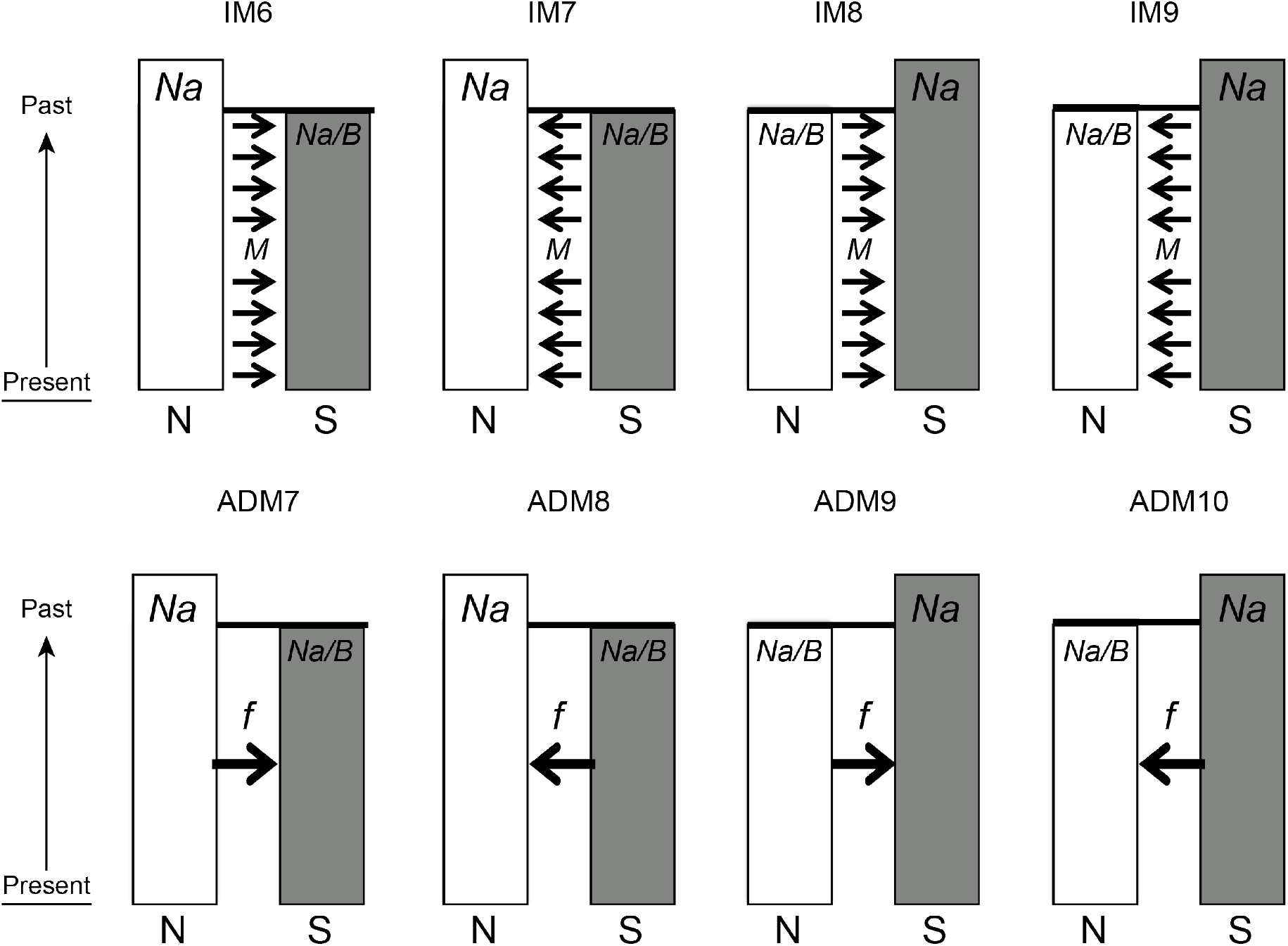
The four population size and gene flow direction topologies fitted for North (N) and South (S) populations of *Pleistodontes nigriventris* in IM (upper row) and ADM models using the blockwise approach. The parameters fitted for IM and ADM models are shown in main text Fig.1. Model names are used to refer to specific models in the main text and in Table 1. For both IM and ADM approaches we compared the models shown with simpler nested models with a single population size (*Na*) and no gene flow (*M*=0 in the IM model, *f*=0 in the ADM model) (models IM 1-5 and ADM 1-6, Table 1).

**Figure S3.**
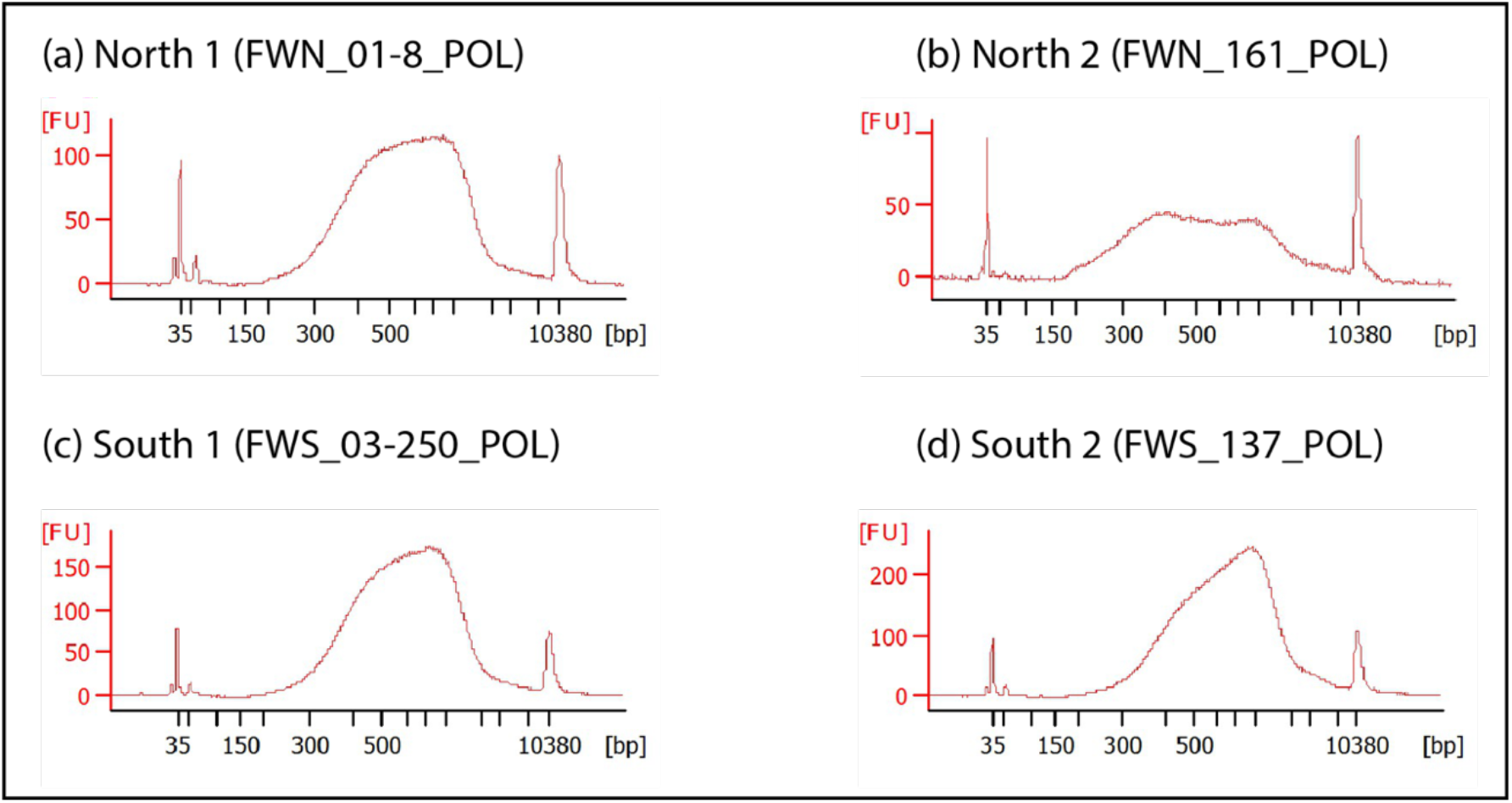
Nextera genomic library fragment size distributions for each of the four male *P. nigriventris*. The x axis in each plot is fragment length in base pairs (bp), and the y axis is relative abundance in fluorescent units (FUs). The distributions were generated from 1μl of each library run on an Agilent 2100 Bioanalyzer using a high sensitivity DNA chip.

**Figure S4.**
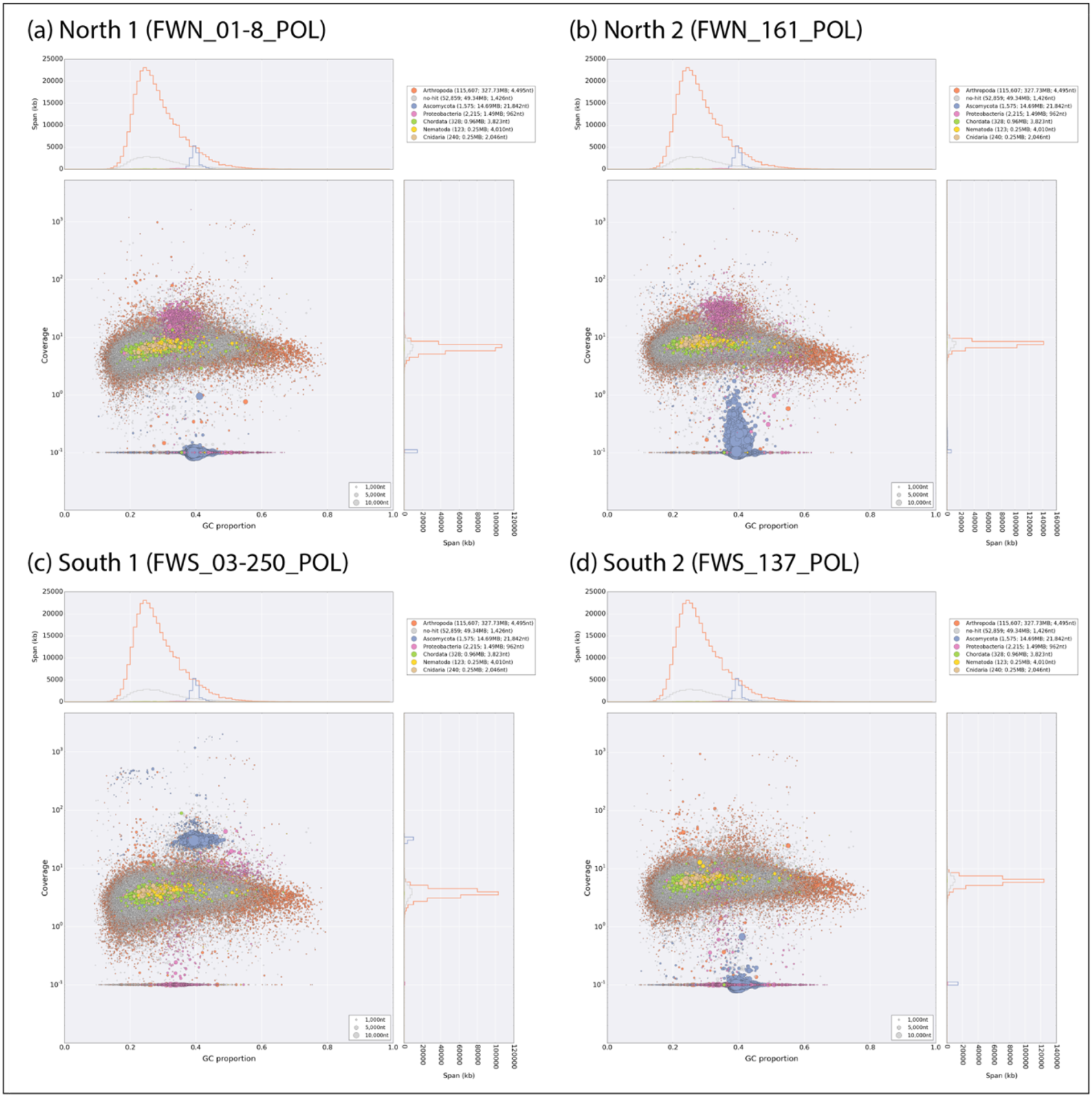
Blobplots for each of the four *P. nigriventris* individuals. Each point shows the GC content (x axis) and coverage (y axis) of a single contig. Points are scaled by contig length and colour-coded by phylum (see key). The distributions above and to the right of each blob plot show the span size for each phylum, e.g. blue blobs represent Ascomycota-designated contigs, pink blobs Proteobacteria-designated contigs (primarily *Wolbachia*). The total number of contigs assigned to each phylum, the total length (span) and the average contig length allocated are given in brackets. Phyla are ranked (top to bottom) in order of declining span.

**Figure S5.**
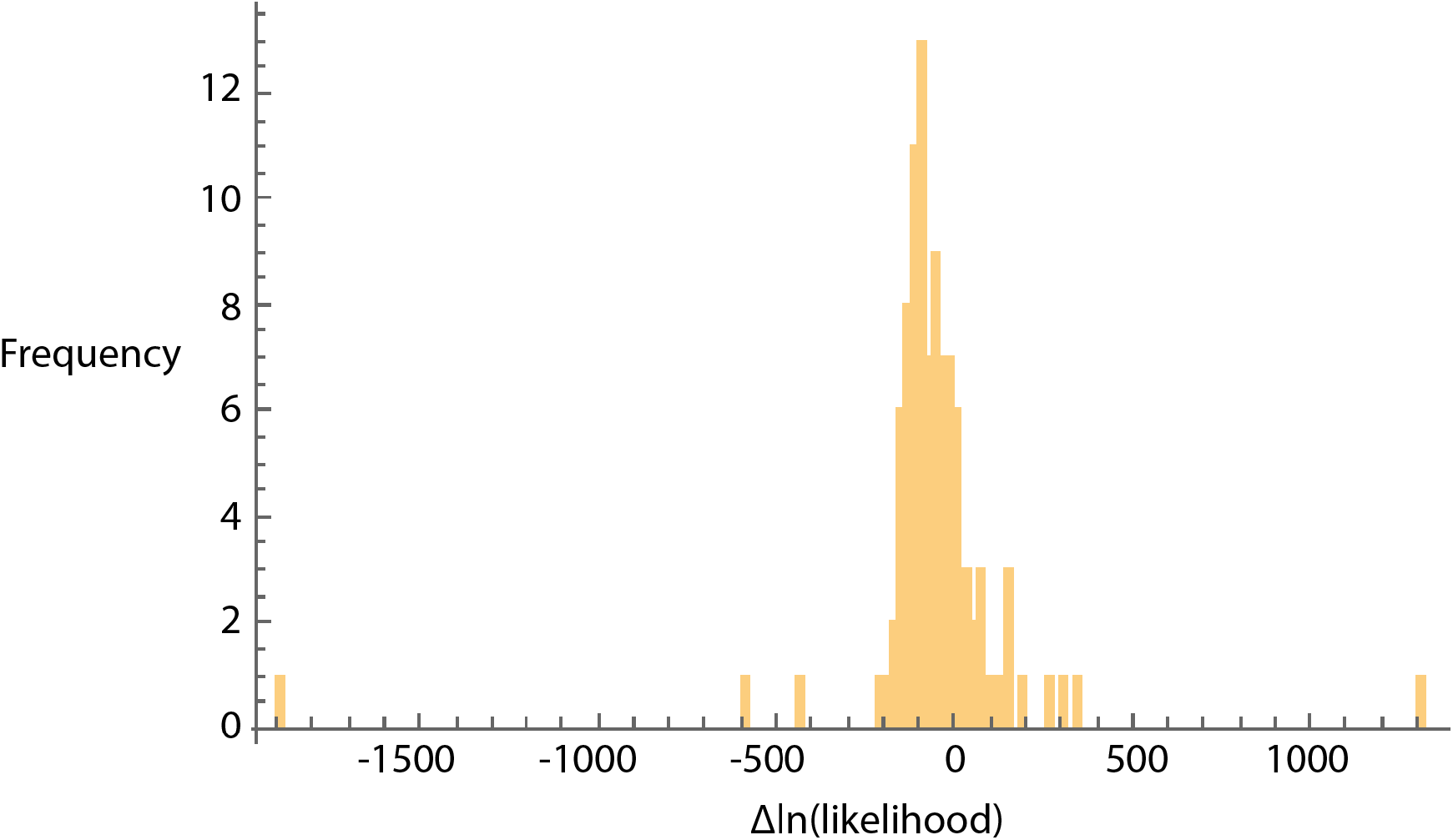
The distribution of relative support (difference in log likelihood, ΔlnL) for the best fitting ADM (ADM10) relative to the IM (IM9) models obtained from datasets simulated under the IM9 model (maximum composite likelihood estimates obtained from the real data). This empirical distribution of ΔlnL values under the null hypothesis (i.e. the simpler IM9 model is true) was used to obtain a critical value of 155.4 at p=0.05.

**Figure S6.**
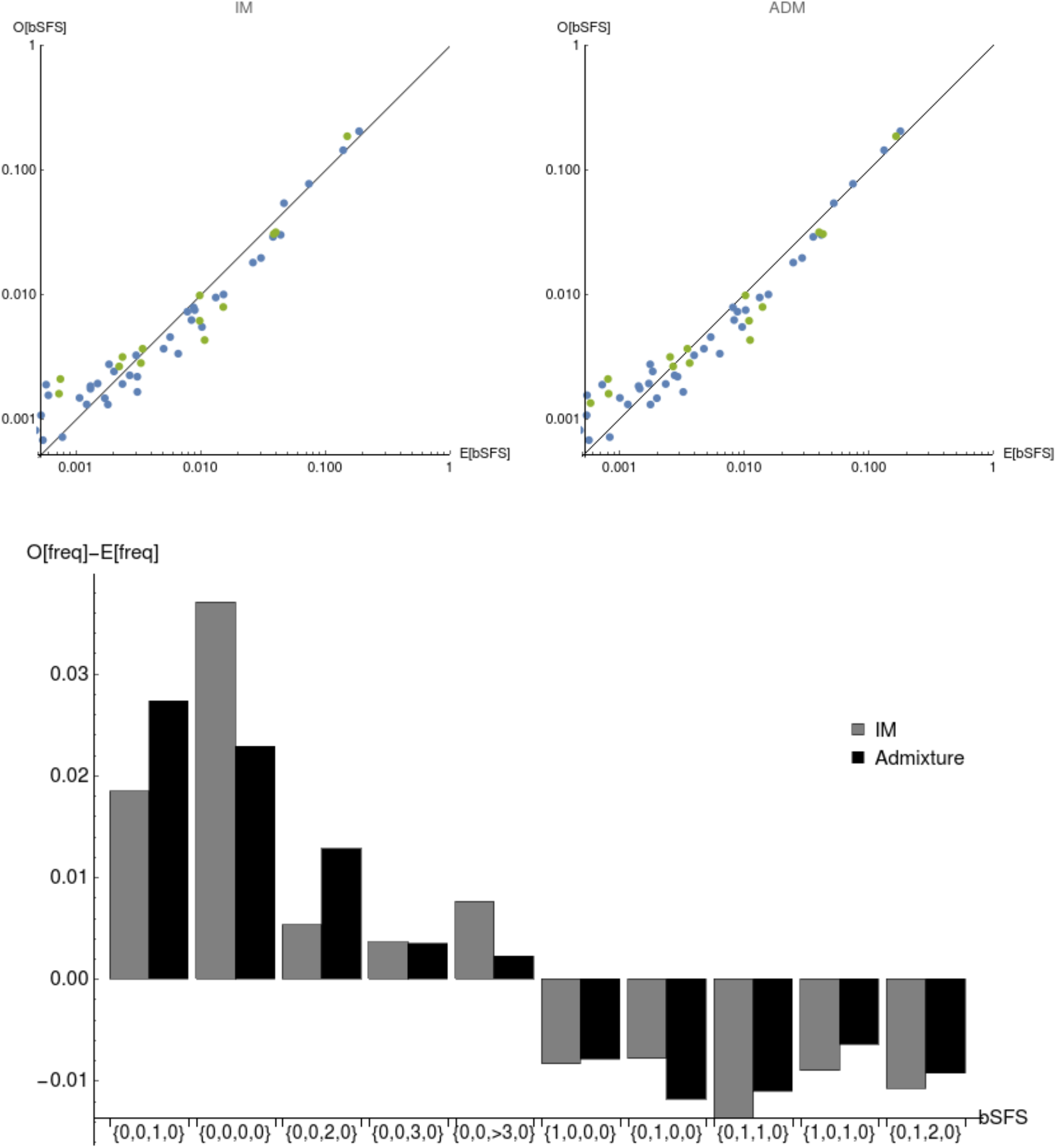
Top) Comparison of the observed frequencies of bSFS configurations with those expected under the best-fitting IM model IM9 (left), and the best fitting ADM model ADM10 (right). A perfect fit to the data would place all points on the diagonal. bSFS configurations without fixed differences are shown in green. Bottom) The residual, i.e. the difference between observed and expected frequency for the 10 most frequent bSFS configurations in the data (are ordered by their frequency in the data from right to left). The ADM model provides a better fit to both monomorphic blocks ({0,0,0,0}, the 2^nd^ most common bSFS configuration in the data) and blocks with a large number of

## References

Aeschbacher, S., Selby, J. P., Willis, J. H. & Coop, G. (2017). Population-genomic inference of the strength and timing of selection against gene flow. Proceedings of the National Academy of Sciences of the USA, 114, 7061–7066. https://doi.org/10.1073/pnas.1616755114

Ahmed, S., Compton, S. G., Butlin, R. K., & Gilmartin P. M. (2009). Wind-borne insects mediate directional pollen transfer between desert fig trees 160 kilometers apart. Proceedings of the National Academy of Sciences of the USA, 106, 20342–20347. https://doi.org/10.1073/pnas.0902213106

Al-beidh, S. (2010). Investigations into stability in the fig/fig-wasp mutualism. Unpublished PhD thesis, Imperial College, London, U.K.

Allendorf, F. W., Hohenlohe, P. A & Luikart, G. (2010). Genomics and the future of conservation genetics. Nature Reviews Genetics, 11, 697–709. https://doi.org/10.1038/nrg2844

Bain, A., Borges, R. M., Chevallier, M. H., Vignes, H., Kobmoo, N., Peng, Y. Q., Cruaud, A., Rasplus, J. Y., Kjellberg, F., & Hossaert-Mckey, M. (2016). Geographic structuring into vicariant species-pairs in a wide-ranging, high-dispersal plant–insect mutualism: the case of *Ficus racemosa* and its pollinating wasps. Evolutionary Ecology, 30, 663–684. https://doi.org/10.1007/s10682-016-9836-5

Bankevich, A., Nurk, S., Antipov, D., Gurevich, A. A., Dvorkin, M., Kulikov, A. S & Pevzner, P. A. (2012). SPAdes: a new genome assembly algorithm and its applications to single cell sequencing. Journal of Computational Biology, 19, 455–477. https://doi.org/10.1089/cmb.2012.0021

Baker, C. H., Graham, G. C., Scott, K. D., Cameron, S. L., Yeates, D. K. & Merritt, D. J. (2008). Distribution and phylogenetic relationships of Australian glow-worms *Arachnocampa* (Diptera, Keroplatidae)’, Molecular Phylogenetics and Evolution, 48, 506–514. https://doi.org/10.1016/j.ympev.2008.04.037

Bell, K. L., Moritz, C., Moussalli, A. & Yeates, D. K. (2007). Comparative phylogeography and speciation of dung beetles from the Australian Wet Tropics rainforest. Molecular Ecology, 16, 4984–4998. https://doi.org/10.1111/j.1365-294X.2007.03533.x

The impact of global selection on local adaptation and reproductive isolation Bisschop, G., Setter, D., Rafajlović, M., Baird, S. J. E. & Lohse, K. (2019). bioRxiv 855320. https://doi.org/10.1101/855320

Bronstein, J. L., Gouyon, P.-H., Gliddon, C., Kjellberg, F. & Michaloud, G. (1990). The ecological consequences of flowering asynchrony in monoecious figs: a simulation study. Ecology, 71, 2145–2156. https://doi.org/10.2307/1938628

Bronstein, J. L. & Hossaert-McKey, M. (1996). Variation in reproductive success within a subtropical fig/pollinator mutualism. Journal of Biogeography, 23, 433–446. https://doi.org/10.1111/j.1365-2699.1996.tb00005.x)

Brown, M., Cooksley, H., Carthew, S M. & Cooper, S. J. B. (2006). Conservation units and phylogeographic structure of an arboreal marsupial, the yellow-bellied glider (*Petaurus australis*). Australian Journal of Zoology, 54, 305–317. https://doi.org/10.1071/ZO06034

Bryant, L. M. & Fuller, S. J. (2014). Pleistocene climate fluctuations influence phylogeographical patterns in *Melomys cervinipes* across the mesic forests of eastern Australia. Journal of Biogeography, 41, 1923–1935. https://doi.org/10.1111/jbi.12341

Bryant, L. M. & Krosch, M. N. (2016). Lines in the land: a review of evidence for eastern Australia’s major biogeographical barriers to closed forest taxa. Biological Journal of the Linnean Society, 119, 238–264. https://doi.org/10.1111/bij.12821

Buchfink, B., Xie, C. & Huson, D. H. (2015). Fast and sensitive protein alignment using DIAMOND. Nature Methods, 12, 59–60. https://doi.org/10.1038/nmeth.3176

Bunnefeld, L., Hearn, J., Stone, G.N. & Lohse, K. (2018). Whole genome data reveal the complex history of a diverse ecological community. Proceedings of the National Academy of Sciences of the United States of America. 115, E6507–E6515. https://doi.org/10.1073/pnas.1800334115

Burke, J. M., Ladiges, P.Y., Batty, E. L., Adams, P. B., Bayly, M. J. & Katinas, L. (2013). Divergent lineages in two species of *Dendrobium* orchids (*D. speciosum* and *D. tetragonum*) correspond to major geographical breaks in eastern Australia. Journal of Biogeography, 40, 2071–2081. https://doi.org/10.1111/jbi.12145

Byrne, M. (2008). Evidence for multiple refugia at different time scales during Pleistocene climatic oscillations in southern Australia inferred from phylogeography. Quaternary Science Reviews, 27, 2576–2585.

Catullo, R. A. & Keogh, J. S. (2014). Aridification drove repeated episodes of diversification between Australian biomes: evidence from a multi-locus phylogeny of Australian toadlets (*Uperoleia*: Myobatrachidae). Molecular Phylogenetics and Evolution, 79, 106–117. https://doi.org/10.1016/j.ympev.2014.06.012

Catullo, R. A., Lanfear, R., Doughty, P. & Keogh, J. S. (2014). The biogeographical boundaries of northern Australia: evidence from ecological niche models and a multi-locus phylogeny of *Uperoleia* toadlets (Anura: Myobatrachidae). Journal of Biogeography, 41, 659–672. https://doi.org/10.1111/jbi.12230

Chapple, D. G., Hoskin, C. J., Chapple, S. N. & Thompson, M. B. (2011). Phylogeographic divergence in the widespread delicate skink (*Lampropholis delicata*) corresponds to dry habitat barriers in eastern Australia. BMC Evolutionary Biology, 11, 191. https://doi.org/10.1186/1471-2148-11-191

Chen, Y., Compton, S. G., Liu, M. & Chen, X. Y. (2012). Fig trees at the northern limit of their range: the distributions of cryptic pollinators indicate multiple glacial refugia. Molecular Ecology, 21, 1687–1701. https://doi.org/10.1111/j.1365-294X.2012.05491.x

Clark, P. U., Hostetler, S. W., Pisias, N. G., Schmittner A. & Meissner, K. J. (2007). Mechanisms for an ∼7-Kyr climate and sea-level oscillation during marine isotope stage 3. In: *Ocean Circulation: Mechanisms and Impacts—Past and Future Changes of Meridional Overturning*. American Geophysical Union. Geophysical Monograph Series, Volume 173. https://doi.org/10.1029/173GM15

Cook, James M. & Rasplus, J.-Y. (2003). Mutualists with attitude: coevolving fig wasps and figs. Trends in Ecology & Evolution, 18, 241–248. https://doi.org/10.1016/S0169-5347(03)00062-4

Cooper, L., Bunnefeld, L., Hearn, J., Cook, J. M., Lohse, K. & Stone, G. N. (2020)a. Phylogeography of Pleistodontes nigriventris from northern and southern populations in Australia. Sequence read data and SPAdes genome assembly in the European Nucleotide Archive, Study PRJEB35527. Available from https://www.ebi.ac.uk/ena/browser/view/PRJEB35527

Cooper, L., Bunnefeld, L., Hearn, J., Cook, J. M., Lohse, K. & Stone, G. N. (2020)b. Phylogeography of *Pleistodontes nigriventris* from northern and southern populations in Australia. Trimmed sequence block data and associated *Mathematica* notebook for blockwise analyses. Available from the Dryad Digital Repository, doi:// (to be added on acceptance).

Costa, R. J. & Wilkinson-Herbots, H. (2017). Inference of gene flow in the process of speciation: an efficient maximum-likelihood method for the isolation-with-initial-migration model. Genetics, 205, 1597–1618. https://doi.org/10.1534/genetics.116.188060.

Darwell, C. T., Al-Beidh, S. & Cook, J. M. (2014). Molecular species delimitation of a symbiotic fig-pollinating wasp species complex reveals extreme deviation from reciprocal partner specificity. BMC Evolutionary Biology, 14, 189. https://doi.org/10.1186/s12862-014-0189-9

DePristo, M. A., Banks, E., Poplin, R., Garimella, K. V., Maguire, J. R., Hartl, C & Daly, M. J. (2011). A framework for variation discovery and genotyping using next-generation DNA sequencing data. Nature Genetics, 43, 491–498. https://doi.org/10.1038/ng.806

Dixon, D. J. (2003). A taxonomic revision of the Australian *Ficus* species in the section Malvanthera (*Ficus* subg. *Urostigma*: Moraceae). Telopea, 10, 125–153. https://doi.org/10.7751/telopea20035611

Dolman, G. & Moritz, C. (2006). A multilocus perspective on refugial isolation and divergence in rainforest skinks (*Carlia*). Evolution, 60, 573–582. https://doi.org/10.1554/05-487.1

Durand E. Y., Patterson N., Reich D., & Slatkin, M. (2011). Testing for ancient admixture between closely related populations. Molecular Biology and Evolution, 28, 2239–2252.. https://doi.org/10.1093/molbev/msr048

Felsenstein, J. 2003. Inferring Phylogenies. Sinauer, 664pp.

Frankham, R. (1995). Effective population size/adult population size ratios in wildlife: a review. Genetics Research, 66, 95–107. https://doi.org/10.1017/S0016672300034455

Frankham, G. J., Handasyde, K. A. & Eldridge, M. D. B (2016). Evolutionary and contemporary responses to habitat fragmentation detected in a mesic zone marsupial, the long-nosed potoroo (*Potorous tridactylus*) in south-eastern Australia. Journal of Biogeography, 43, 653–665. https://doi.org/10.1111/jbi.12659

Fuentes-Pardo, A. P. & Ruzzante, D. E. (2017). Whole-genome sequencing approaches for conservation biology: Advantages, limitations and practical recommendations. Molecular Ecology, 26, 5369–5406. https://doi.org/10.1111/mec.14264

García-Alcalde, F., Okonechnikov, K., Carbonell, J., Cruz, L. M., Götz, S., Tarazona, S., Dopazo, J., Meyer, T. F. & Conesa, A. (2012). Qualimap: evaluating next-generation sequencing alignment data. Bioinformatics, 28, 2678–2679. https://doi.org/10.1093/bioinformatics/bts503

Greeff, J. M., van Vuuren, G. J. J., Kryger, P. & Moore, J. C. (2009). Outbreeding and possibly inbreeding depression in a pollinating fig wasp with a mixed mating system. Heredity, 102, 349–356. https://doi.org/10.1038/hdy.2009.2

Haine, E. R., & Cook, J. M. (2005). Convergent incidences of *Wolbachia* infection in fig wasp communities from two continents. Proceedings of the Royal Society of London, Series B Biological Sciences, 272, 421–429. https://doi.org/10.1098/rspb.2004.2956

Hare, M. P. (2001). Prospects for nuclear gene phylogeography. Trends in Ecology & Evolution, 16, 700–706. https://doi.org/10.1016/S0169-5347(01)02326-6

Harrison, R. D. (2000). Repercussions of El Niño: drought causes extinction and the breakdown of mutualism in Borneo. Proceedings of the Royal Society of London, Series B Biological Sciences, 267, 911–915. https://doi.org/10.1098/rspb.2000.1089

Harrison, R. D. (2003). Fig wasp dispersal and the stability of a keystone plant resource in Borneo. Proceedings of the Royal Society of London, Series B Biological Sciences, 270, Suppl 1: S76–9.

Harrison, R. D. (2005). Figs and the diversity of tropical rainforests. Bioscience, 55, 1053–1064. https://doi.org/10.1098/rsbl.2003.0018

Harrison, R. D. & Rasplus, J.-Y. (2006). Dispersal of fig pollinators in Asian tropical rain forests. Journal of Tropical Ecology, 22, 631–639. https://doi.org/10.1017/S0266467406003488

Hearn, J., Stone, G. N., McInnes, L., Nicholls, J. A., Barton, N. H. & Lohse, K. (2014). Likelihood-based inference of population history from low coverage de novo genome assemblies. Molecular Ecology, 23, 198–211. https://doi.org/10.1111/mec.12578.

Hedrick, P. W. (2013). Adaptive introgression in animals: examples and comparison to new mutation and standing variation as sources of adaptive variation. Molecular Ecology, 22, 4606–4618. https://doi.org/10.1111/mec.12415

Hey, J. & Nielsen, R. (2004). Multilocus methods for estimating population sizes, migration rates and divergence time, with applications to the divergence of *Drosophila pseudoobscura* and *D. persimilis*. Genetics, 167, 747–60. https://doi.org/10.1534/genetics.103.024182.

Hocknull, S. A., Zhao, J.-X., Feng, Y.-X. & Webb, G. E. (2007). Responses of Quaternary rainforest vertebrates to climate change in Australia. Earth and Planetary Science Letters, 264, 317–331. https://doi.org/10.1016/j.epsl.2007.10.004

Jennings, W. B. & Edwards, S. V. (2005). Speciational history of Australian grass finches (*Poephila*) inferred from thirty gene trees. Evolution, 59, 2033–2047. https://doi.org/10.1111/j.0014-3820.2005.tb01072.x

Jermiin, L. S. & Crozier, R. H. (1994). The cytochrome b region in the mitochondrial DNA of the ant *Tetraponera rufoniger*: sequence divergence in Hymenoptera may be associated with nucleotide content. Journal of Molecular Evolution, 38, 282–294. https://doi.org/10.1007/BF00176090

Jesus, F. F., Wilkins, J. F., Solferini, V. N. & Wakeley, J. (2006). Expected coalescence times and segregating sites in a model of glacial cycles. Genetic and Molecular Research, 5, 466–474.

Jia, X. C., Yao, J. Y., Chen, Y. Z., Cook, J. M & Crozier, R. H. (2008). The phenology and potential for self-pollination of two Australian monoecious fig species. Symbiosis, 45, 91–96.

Keightley, P. D., Ness, R. W., Halligan, D. L. & Haddrill, P. R. (2014). Estimation of the spontaneous mutation rate per nucleotide site in a *Drosophila melanogaster* full-sib family. Genetics, 196, 313–320. https://doi.org/10.1534/genetics.113.158758

Keightley, P. D., Pinharanda, A., Ness, R. W., Simpson, F., Dasmahapatra, K. K., Mallet, J., Davey, J. W. & Jiggins C. D. (2015). Estimation of the spontaneous mutation rate in *Heliconius melpomene*. Molecular Biology and Evolution, 32, 239–243. https://doi.org/10.1093/molbev/msu302

Kelleher, J., Etheridge, A. M. & McVean, G. (2016). Efficient coalescent simulation and genealogical analysis for large sample sizes. PLoS Computational Biology, 12, e1004842. https://doi.org/10.1371/journal.pcbi.1004842 https://doi.org/10.1371/journal.pcbi.1004842

Kershaw, A. P. (1994). Pleistocene vegetation of the humid tropics of northeastern Queensland, Australia. Paleoclimatology, Paleogeography, Paleoecology, 109, 399–412. https://doi.org/10.1016/0031-0182(94)90188-0 https://doi.org/10.1016/0031-0182(94)90188-0

Kimura, M. & Crow, J. F. (1963). The measurement of effective population number. Evolution, 17, 279–288. https://doi.org/10.1111/j.1558-5646.1963.tb03281.x

Kobmoo, N., Hossaert-McKey, M., Rasplus, J.-Y. & Kjellberg, F. (2010). *Ficus racemosa* is pollinated by a single population of a single agaonid wasp species in continental South-East Asia. Molecular Ecology, 19, 2700–2712. https://doi.org/10.1111/j.1365-294X.2010.04654.x

Kumar, S., Jones, M., Koutsovoulos, G., Clarke, M. & Blaxter, M. (2013). Blobology: exploring raw genome data for contaminants, symbionts and parasites using taxon-annotated GC-coverage plots’, Frontiers in Genetics, 4, 237. https://doi.org/10.3389/fgene.2013.00237

Langmead, B., & Salzberg, S. L. (2012). Fast gapped-read alignment with Bowtie 2. Nature Methods, 9, 357–359. https://doi.org/10.1038/nmeth.1923

Lanier, H. C., & Knowles, L. L.(2012). Is recombination a problem for species-tree analyses? Systematic Biology, 61, 691–701. https://doi.org/10.1093/sysbio/syr128

Leffler, E. M., Bullaughey, K., Matute, D. R., Meyer, W. K., Ségurel, L., Venkat, A., Andolfatto, P. & Przeworski, M. (2012). Revisiting an old riddle: what determines genetic diversity levels within species? PLoS Biology, 10(9), e1001388. https://doi.org/10.1371/journal.pbio.1001388

Lewontin, R. C. (1974). The Genetic Basis of Evolutionary Change. Columbia University Press, New York. 346 pp.

Li, H. (2011). A statistical framework for SNP calling, mutation discovery, association mapping and population genetical parameter estimation from sequencing data. Bioinformatics, 27, 2987–2993. https://doi.org/10.1093/bioinformatics/btr509

Li, H. & Durbin, R. (2009). Fast and accurate short read alignment with Burrows-Wheeler transform. Bioinformatics, 25, 1754–1760. https://doi.org/10.1093/bioinformatics/btp324

Li, H. & Durbin, R. (2011). Inference of human population history from individual whole-genome sequences. Nature, 475, 493–496. https://doi.org/10.1038/nature10231

Li, H., Handsaker, B., Wysoker, A., Fennell, T., Ruan, J., Homer, N., Marth, G., Abecasis, G., Durbin, R. & Subgroup Genome Project Data Processing. (2009). The Sequence Alignment/Map format and SAMtools. Bioinformatics, 25, 2078–2079. https://doi.org/10.1093/bioinformatics/btp352

Liu, M., Zhao, R., Chen, Y., Zhang, J., Compton, S. G. & Chen, X. Y. (2014). Competitive exclusion among fig wasps achieved via entrainment of host plant flowering phenology. PLoS ONE 9(5): e97783. doi:10.1371/journal.pone.0097783

Liu, M., Compton, S. G., Peng, F. E., Zhang, J. & Chen, X. Y. (2015). Movements of genes between populations: are pollinators more effective at transferring their own or plant genetic markers?’, Proceedings of the Royal Society of London, Series B Biological Sciences, 282, 20150290. https://doi.org/10.1098/rspb.2015.0290

Lohse, K., Chmelik, M., Martin, S. H. & Barton, N. H. (2016). Efficient strategies for calculating blockwise likelihoods under the coalescent. Genetics, 202, 775–786. https://doi.org/10.1534/genetics.115.183814

Lohse, K. & Frantz, L. A. F. (2014). Neandertal admixture in Eurasia confirmed by maximum-likelihood analysis of three genomes. Genetics, 196, 1241–1251. https://doi.org/10.1534/genetics.114.162396

Lohse, K., Harrison, R. J. & Barton, N. H. (2011). A general method for calculating likelihoods under the coalescent process. Genetics, 189, 977–987. https://doi.org/10.1534/genetics.111.129569

Lopez-Vaamonde, C., Rasplus, J.Y., Weiblen, G.D., and Cook, J.M. (2001) Molecular phylogenies of fig wasps: Partial cocladogenesis of pollinators and parasites. Molecular Phylogenetics and Evolution, 21, 55–71. https://doi.org/10.1006/mpev.2001.0993

Lopez-Vaamonde, C., Dixon, D. J., Cook, J. M. & Rasplus, J.-Y. (2002). Revision of the Australian species of *Pleistodontes* (Hymenoptera: Agaonidae) fig-pollinating wasps and their host-plant associations. Zoological Journal of the Linnean Society, 136, 637–683. https://doi.org/10.1046/j.1096-3642.2002.00040.x

Macqueen, P., Seddon, J. M., Austin, J. J., Hamilton, S. & Goldizen, A. W. (2010). Phylogenetics of the pademelons (Macropodidae: *Thylogale*) and historical biogeography of the Australo-Papuan region. Molecular Phylogenetics and Evolution, 57, 1134–1148. https://doi.org/10.1016/j.ympev.2010.08.010

Macqueen, P., Seddon, J. M. & Goldizen, A. W. (2012). Effects of historical forest contraction on the phylogeographic structure of Australo-Papuan populations of the red-legged pademelon (Macropodidae: *Thylogale stigmatica*). Austral Ecology, 37, 479–490. https://doi.org/10.1111/j.1442-9993.2011.02309.x

Maldonado, S. P., Melville, J., Peterson, G. N. L. & Sumner, J. (2012). Human-induced versus historical habitat shifts: identifying the processes that shaped the genetic structure of the threatened grassland legless lizard, *Delma impar*. Conservation Genetics, 13, 1329-1342. https://doi.org/10.1007/s10592-012-0377-3

Male, T. D. & Roberts, G. E. (2005). Host associations of the strangler fig *Ficus watkinsiana* in a subtropical Queensland rain forest. Austral Ecology, 30, 229–236. https://doi.org/10.1111/j.1442-9993.2005.01442.x

Markgraf, V., McGlone, M. & Hope, G. (1995). Neogene paleoenvironmental and paleoclimatic change in southern temperate ecosystems — a southern perspective. Trends in Ecology & Evolution, 10, 143–147. https://doi.org/10.1016/S0169-5347(00)89023-0

Martin, H. A. (2006). Cenozoic climatic change and the development of the arid vegetation in Australia. Journal of Arid Environments, 66, 533–563. https://doi.org/10.1016/j.jaridenv.2006.01.009

Martin, M. (2011). Cutadapt removes adapter sequences from high-throughput sequencing reads. EMBnet.journal, 17, No 1: Next Generation Sequencing Data Analysis. https://doi.org/https://doi.org/10.14806/ej.17.1.200

Martin, S. H., Dasmahapatra, K. K., Nadeau, N. J., Salazar, C., Walters, J. R., Simpson, F., Blaxter, M., Manica, A., Mallet, J. & Jiggins, C. D. (2013). Genome-wide evidence for speciation with gene flow in *Heliconius* butterflies. Genome Research, 23, 1817–1828. https://doi.org/10.1101/gr.159426.113

McGuigan, K., McDonald, K., Parris, K. & Moritz, C. (1998). Mitochondrial DNA diversity and historical biogeography of a wet forest-restricted frog (*Litoria pearsoniana*) from mid-east Australia. Molecular Ecology, 7, 175–186. https://doi.org/10.1046/j.1365-294x.1998.00329.x

McKenna, A., Hanna, M., Banks, E., Sivachenko, A., Cibulskis, K., Kernytsky, A., Garimella, K., Altshuler, D., Gabriel, S., Daly, M. & DePristo, M. A. (2010). The Genome Analysis Toolkit: a MapReduce framework for analyzing next generation DNA sequencing data. Genome Research, 20, 1297–1303. https://doi.org/10.1101/gr.107524.110

Molbo, D., Machado, C. A., Herre, E. A. & Keller, L. (2004). Inbreeding and population structure in two pairs of cryptic fig wasp species. Molecular Ecology, 13, 1613–1623. https://doi.org/10.1111/j.1365-294X.2004.02158.x

Nazareno, A. G., Alzate-Marin, A. L. & Pereira, R. A. S. (2013). Dioecy, more than monoecy, affects plant spatial genetic structure: the case study of *Ficus*. Ecology and Evolution, 3, 3495–3508. https://doi.org/doi:10.1002/ece3.739

Nicholls, J. A., & Austin, J. J. (2005). Phylogeography of an east Australian wet forest bird, the satin bowerbird (*Ptilonorhynchus violaceus*), derived from mtDNA, and its relationship to morphology. Molecular Ecology, 14, 1485–1496. https://doi.org/10.1111/j.1365-294X.2005.02544.x

Nielsen, R. & Wakeley, J. (2001). Distinguishing migration from isolation: a Markov Chain Monte Carlo approach. Genetics, 158, 885–896.

Niu, L.-H., Song, X.-F., He, S. M., Zhang, P., Wang, N. X., Li, Y. & Huang, D. W. (2015). New insights into the fungal community from the raw genomic sequence data of fig wasp *Ceratosolen solmsi*. BMC Microbiology, 15, 27. https://doi.org/10.1186/s12866-015-0370-3

Nürnberger, B., Lohse, K., Fijarczyk, A., Szymura, J. M. & Blaxter, M. L. (2016). Para-allopatry in hybridizing fire-bellied toads (*Bombina bombina* and *B. variegata*): inference from transcriptome-wide coalescence analyses. Evolution, 70, 1803–1818. https://doi.org/10.1111/evo.12978

Okonechnikov, K., Conesa, A. & García-Alcalde, F (2016). Qualimap 2: advanced multi-sample quality control for high-throughput sequencing data. Bioinformatics, 32, 292–294. https://doi.org/10.1093/bioinformatics/btv566

Oswald, J. A., Overcast, I., Mauck, W. M., Andersen, M. J. & Smith, B. T. (2017). Isolation with asymmetric gene flow during the nonsynchronous divergence of dry forest birds. Molecular Ecology, 26, 1386–1400. https://doi.org/10.1111/mec.14013

Palstra, F. P. & Ruzzante, D. E. (2008). Genetic estimates of contemporary effective population size: what can they tell us about the importance of genetic stochasticity for wild population persistence? Molecuar Ecology, 17, 3428-3447. https://doi.org/10.1111/j.1365-294X.2008.03842.x

Pope, L. C., Estoup, A. & Moritz, C. (2000). Phylogeography and population structure of an ecotonal marsupial, *Bettongia tropica*, determined using mtDNA and microsatellites. Molecular Ecology, 9, 2041–2053. https://doi.org/10.1046/j.1365-294x.2000.01110.x

Pope, L. C., Storch, D., Adams, M., Moritz, C. & Gordon, G. (2001). A phylogeny for the genus *Isoodon* and a range expansion *for I. obesulus peninsulae* based on mtDNA control region and morphology. Australian Journal of Zoology, 49, 411–434. https://doi.org/10.1071/ZO00060

Price, A. L., Jones, N. C. & Pevzner, P. A. (2005). *De novo* identification of repeat families in large genomes. Bioinformatics, 21 Suppl 1, i351-i358. https://doi.org/10.1093/bioinformatics/bti1018

Quinlan, A. R. & Hall, I. M. (2010). BEDTools: a flexible suite of utilities for comparing genomic features, Bioinformatics, 26, 841–842. https://doi.org/10.1093/bioinformatics/btq033

Ringbauer, H., Kolesnikov, A., Field, D. L. & Barton, N. H. (2018). Estimating barriers to gene flow from distorted isolation-by-distance patterns. Genetics, 208, 1231–1245. https://doi.org/10.1534/genetics.117.300638

Rix, M. G. & Harvey, M. S. (2012). Phylogeny and historical biogeography of ancient assassin spiders (Araneae: Archaeidae) in the Australian mesic zone: evidence for Miocene speciation within Tertiary refugia. Molecular Phylogenetics and Evolution, 62, 375–396. https://doi.org/10.1016/j.ympev.2011.10.009

Rodriguez, L. J., Bain, A., Chou, L.-S., Conchou, L., Cruaud, A., Gonzales, R., Hossaert-McKey, M., Rasplus, J.-Y., Tzeng, H.-Y. & Kjellberg, F. (2017). Diversification and spatial structuring in the mutualism between *Ficus septica* and its pollinating wasps in insular South East Asia. BMC Evolutionary Biology, 17, 207. https://doi.org/10.1186/s12862-017-1034-8

Romiguier, J., Gayral, P., Ballenghien, M., Bernard, A., Cahais, V., Chenuil, A., Chiari, Y., Dernat, R., Duret, L., Faivre, N., Loire, E., Lourenco, J. M., Nabholz, B., Roux, C., Tsagkogeorga, G., Weber, A. A.-T., Weinert, L. A., Belkhir, K., Bierne, N. Glémin S. & Galtier, N. (2014). Comparative population genomics in animals uncovers the determinants of genetic diversity. Nature, 515, 261–263. https://doi.org/10.1038/nature13685

Rønsted, N., Weiblen, G. D., Savolainen, V. & Cook, J. M. (2008). Phylogeny, biogeography, and ecology of *Ficus* section Malvanthera (Moraceae). Molecular Phylogenetics and Evolution, 48, 12–22. https://doi.org/10.1016/j.ympev.2008.04.005

Schiffer, M., Kennington, W. J., Hoffmann, A. A. & Blacket, M. J. (2007). Lack of genetic structure among ecologically adapted populations of an Australian rainforest *Drosophila* species as indicated by microsatellite markers and mitochondrial DNA sequences. Molecular Ecology, 16, 1687–700. . https://doi.org/10.1111/j.1365-294X.2006.03200.x

Schäuble, C. S., & Moritz, C. (2001). Comparative phylogeography of two open forest frogs from eastern Australia. Biological Journal of the Linnean Society, 74, 157–70. https://doi.org/10.1111/j.1095-8312.2001.tb01384.x

Schneider, C. J., Cunningham, M. & Moritz, C. (1998). Comparative phylogeography and the history of endemic vertebrates in the Wet Tropics rainforests of Australia. Molecular Ecology, 7, 487–498. https://doi.org/10.1046/j.1365-294x.1998.00334.x

Segar, S. T., Dunn, D. W., Darwell, C. T. & Cook, J. M. (2014). How to be a fig wasp down under: the diversity and structure of an Australian fig wasp community. Acta Oecologica, 57, 17–27. https://doi.org/10.1016/j.actao.2013.03.014

Siddall, M., Rohling, E. J., Thompson, W. G. & Waelbroeck, C. (2008). Marine isotope stage 3 sea level fluctuations: data synthesis and new outlook. Reviews of Geophysics, 46. https://doi.org/10.1029/2007RG000226

Simao, F. A., Waterhouse, R. M., Ioannidis, P., Kriventseva, E. V. & Zdobnov, E. M. (2015). BUSCO: assessing genome assembly and annotation completeness with single-copy orthologs. Bioinformatics, 31, 3210–3212. https://doi.org/10.1093/bioinformatics/btv351

Smissen, P. J., Melville, J., Sumner, J. & Jessop, T. S. (2013). Mountain barriers and river conduits: phylogeographical structure in a large, mobile lizard (Varanidae: *Varanus varius*) from eastern Australia. Journal of Biogeography, 40, 1729–1740. https://doi.org/10.1111/jbi.12128

Smit, A. F. A., Hubley, R. & Green, P. (2013-2015). RepeatMasker Open-4.0. 2013-2015 <http://www.repeatmasker.org>.<http://www.repeatmasker.org>

Sousa, V. & Hey, J. (2013). Understanding the origin of species with genome-scale data: modelling gene flow. Nature Reviews Genetics, 14, 404–414. https://doi.org/10.1038/nrg3446

Stone, G. N., White, S. C., Csóka, G., Melika, G., Mutun, S., Pénzes, Z., Sadeghi, S. E., Schönrogge, K., Tavakoli, M. & Nicholls, J. A. (2017). Tournament ABC analysis of the western palaearctic population history of an oak gallwasp, *Synergus umbraculus*. Molecular Ecology, 26, 6685–6703. doi: 10.1111/mec.14372

Strasburg, J. L., & Rieseberg, L. H. (2010). How robust are “Isolation with Migration” analyses to violations of the IM model? A simulation Study. Molecular Biology and Evolution, 27, 297–310. https://doi.org/10.1093/molbev/msp233

Stuart-Fox, D. M., Schneider, C. J., Moritz, C. & Couper, P. J. (2001). Comparative phylogeography of three rainforest restricted lizards from mid-east Queensland. Australian Journal of Zoology, 49, 119–127. https://doi.org/10.1071/ZO00092

Stumpf, M. P. & McVean, G. A. (2003). Estimating recombination rates from population-genetic data. Nature Reviews Genetics, 4, 959–68. https://doi.org/10.1038/nrg1227

Sutton, T.L., DeGabriel, J. L., Riegler, M. & Cook, J.M. (2018). A temperate pollinator with high thermal tolerance is still susceptible to heat events predicted under future climate change. Ecological Entomology, 43, 506–512. https://doi.org/10.1111/een.12528

Sutton, T. L., Riegler, M. & Cook, J. M. (2016). One step ahead: a parasitoid disperses farther and forms a wider geographic population than its fig wasp host. Molecular Ecology, 25, 882–94. https://doi.org/10.1111/mec.13445

Tian, E., Nason, J. D., Machado, C. A., Zheng, L., Yu, H. & Kjellberg, F. (2015). Lack of genetic isolation by distance, similar genetic structuring but different demographic histories in a fig-pollinating wasp mutualism. Molecular Ecology, 24, 5976–5991. https://doi.org/10.1111/mec.13438

Treangen, T. J. & Salzberg, S. L. (2011). Repetitive DNA and next-generation sequencing: computational challenges and solutions. Nature Reviews Genetics, 13, 36–46. https://doi.org/10.1038/nrg3117

Van der Auwera, G. A., Carneiro, M. O., Hartl, C., Poplin, R., Del Angel, G., Levy-Moonshine, A., Jordan, T., Shakir, K., Roazen, D., Thibault, J., Banks, E., Garimella, K., Altshuler, D., Gabriel, S. & DePristo M, (2013). From FastQ data to high confidence variant calls: the Genome Analysis Toolkit best practices pipeline. Current Protocols in Bioinformatics, 43, 11.10.1–33. https://doi.org/10.1002/0471250953.bi1110s43.

Van Meerbeeck, C. J., Renssen, H. & Roche, D. M. (2009). How did Marine Isotope Stage 3 and Last Glacial Maximum climates differ? – Perspectives from equilibrium simulations. Climate of the Past, 5, 33–51. https://doi.org/10.5194/cp-5-33-2009

Wachi, N., Kusumi, J., Tzeng, H. Y. & Su, Z. H. (2016). Genome-wide sequence data suggest the possibility of pollinator sharing by host shift in dioecious figs (Moraceae, *Ficus*). Molecular Ecology, 25, 5732–5746. https://doi.org/10.1111/mec.13876

Wall, J. D. (2003). Estimating ancestral population sizes and divergence times. Genetics, 163, 395–404.

Walton, W., Stone, G. N. & Lohse, K. (2020). Diverse demographic histories in a guild of hymenopteran parasitoids. Molecular Ecology, (in review). Add information or delete as required.

Wang, Z. & Liu, K. J. (2016). A performance study of the impact of recombination on species tree analysis. BMC Genomics, 17, 785. https://doi.org/10.1186/s12864-016-3104-5

Ware, A. B. & Compton, S. G. (1994)a. Dispersal of adult female fig wasps. 1. Arrivals and departures. Entomologia Experimentalis et Applicata, 73, 221–229. https://doi.org/10.1111/j.1570-7458.1994.tb01859.x

Ware, A. B. & Compton, S. G. (1994)b. Dispersal of adult female fig wasps. 2. Movements between trees. Entomologia Experimentalis et Applicata, 73, 231–238. https://doi.org/10.1111/j.1570-7458.1994.tb01860.x

Weber, L. C., Van der Wal, J., Schmidt, S., McDonald, W. J. F., Shoo, L. P. & Ladiges, P. (2014). Patterns of rain forest plant endemism in subtropical Australia relate to stable mesic refugia and species dispersal limitations. Journal of Biogeography, 41, 222–38. https://doi.org/10.1111/jbi.12219

Werren, J. H., Richards, S., Desjardins, C. A., Niehuis, O., Gadau, J. & Colbourne, J. K. (2010). Functional and evolutionary insights from the genomes of three parasitoid *Nasonia* species. Science, 327, 343–348. https://doi.org/10.1126/science.1178028

Xiao, J. H., Yue, Z., Jia, L. Y., Yang, X. H., Niu, L. H., Wang, Z., … Huang, D. W. (2013). Obligate mutualism within a host drives the extreme specialization of a fig wasp genome. Genome Biology, 14, R141. https://doi.org/10.1186/gb-2013-14-12-r141

Yang, L. Y., Machado, C. A., Dang, X. D., Peng, Y. Q., Yang, D. R., Zhang, D. Y. & Liao, W. J. (2015). The incidence and pattern of co-pollinator diversification in dioecious and monoecious figs. Evolution, 69, 294–304. https://doi.org/10.1111/evo.12584

Yu, H., Tian, E. W., Zheng, L. N., Deng, X. X., Cheng, Y. F., Chen, L. F & Kjellberg, F. (2019) Multiple parapatric pollinators have radiated across a continental fig tree displaying clinal genetic variation. Molecular Ecology, 28, 2391-2405. https://doi.org/10.1111/mec.15046

Zerbino, D. R. & Birney, E. (2008). Velvet: algorithms for *de novo* short read assembly using de Bruijn graphs. Genome Research, 18, 821–9. https://doi.org/10.1101/gr.074492.107

Zhang, J., Kobert, K., Flouri, T. & Stamatakis, A. (2014). PEAR: a fast and accurate Illumina Paired-End reAd mergeR. Bioinformatics, 30, 614–620. https://doi.org/10.1093/bioinformatics/btt593

## References

Parra, G., Bradnam, K. & Korf, I. (2007). CEGMA: a pipeline to accurately annotate core genes in eukaryotic genomes. Bioinformatics, 23, 1061–1067.

Parra, G., Bradnam, K., Ning, Z., Keane, T. & Korf, I. (2009). Assessing the gene space in draft genomes. Nucleic Acids Research, 37, 289–297.

## Reference

Pedersen, B. S. & Quinlan, A. R. (2018). Mosdepth: quick coverage calculation for genomes and exomes. Bioinformatics, 34(5): 867–868. doi: 10.1093/bioinformatics/btx699

